# Evolutionary processes shaping diversity across the *Homo* lineage

**DOI:** 10.1101/136507

**Authors:** Lauren Schroeder, Rebecca Rogers Ackermann

## Abstract

Recent fossil finds have highlighted extensive morphological diversity within our genus, *Homo*, and the co-existence of a number of species. However, little is known about the evolutionary processes responsible for producing this diversity. Understanding the action of these processes can provide insight into how and why our lineage evolved and diversified. Here, we examine cranial and mandibular variation and diversification from the earliest emergence of our genus at 2.8 Ma until the Late Pleistocene (0.126-0.0117 Ma), using statistical tests developed from quantitative genetics theory to evaluate whether stochastic (genetic drift) versus non-stochastic (selection) processes were responsible for the observed variation. Results show that random processes can account for species diversification for most traits, including neurocranial diversification, and across all time periods. Where selection was found to shape diversification, we show that: 1) adaptation was important in the earliest migration of *Homo* out of Africa; 2) selection played a role in shaping mandibular and maxillary diversity among *Homo* groups, possibly due to dietary differences; and 3) *Homo rudolfensis* is adaptively different from other early *Homo* taxa, including the earliest known *Homo* specimen. These results show that genetic drift, and likely small population sizes, were important factors shaping the evolution of *Homo* and many of its novel traits, but that selection played an essential role in driving adaptation to new contexts.

## Introduction

Our genus is characterized by a significant amount of morphological diversity, a phenomenon at the heart of the longstanding debate surrounding the origin and evolution of *Homo* (see Wood, 1992; Wood and Baker, 2011; Antón et al., 2014). Since the announcement of the fossil remains of *Homo habilis* from Olduvai Gorge over fifty years ago (Leakey et al., 1964) the focus in paleoanthropology has been on trying to tease apart inter- and intra-specific variation within *Homo* to answer questions relating to taxonomic relationships between species (e.g. Miller, 1991, 2000; Wood, 1993; Kramer et al., 1995; Lieberman et al., 1996). However, an ever growing fossil record and an exceedingly variable genus make this a complicated undertaking. Recent fossil finds, such as the geographically extreme and highly variable sample of early *Homo* from Dmanisi, Georgia (∼1.8 Ma; Lordkipanidze et al, 2013), the oldest known specimen of *Homo* from Ledi-Geraru, Ethiopia (∼2.8 Ma; Villmoare et al., 2015), and the derived but small-brained *Homo naledi* from the Dinaledi cave, South Africa (236-335 ka; Berger et al., 2015; Dirks et al., 2017), once again prove that not only is *Homo* diverse and at times mosaic in nature, but that our previous attempts to define and confine *Homo* to a specific suite of characters at a specific time and place are no longer appropriate.

What drives such a degree of diversification and innovation? Unfortunately our understanding of the underlying evolutionary processes acting on *Homo* is limited. Explanations for major transitions in human evolution have tended to focus on adaptive evolutionary scenarios, specifically directional selection acting on a given trait. As one example, the emergence of the genus *Homo,* and its associated big brain and tools (but see Harmand et al., 2015), has been interpreted as an adaptive response to substantial environmental change in Africa ca. 2.5 Ma (Vrba, 1985, 1995, 1996, 2007; Cerling, 1992; Stanley, 1992; deMenocal, 1995; Reed, 1997; Bobe and Behrensmeyer, 2004; Wynn, 2004). However, the limited work examining the evolutionary processes during this transition suggests otherwise, pointing to drift as a major player (Schroeder et al., 2014). Adding additional complexity to this picture, the emerging notion of a highly variable genus, with unanticipated traits such as wide ranges of brain size within species (Spoor et al., 2015), and the re-evolution of small brains in multiple contexts (Brown et al., 2004; Berger et al., 2015), challenge a linear notion of the emergence of *Homo*-like morphology. Instead these data support the idea that the evolution of *Homo* may have been characterized by multiple lineages, and defined by evolutionary innovation and experimentation (Antón et al., 2014). In such a scenario, what we identify as *Homo*-like morphology could have evolved repeatedly, in different contexts or at different times. Yet our understanding of how and why this diversity came to be remains largely unknown. Interestingly, the recent suggestion that habitat fragmentation, as a consequence of major environmental change, was the potential driving force behind the diversification of early *Homo* (Antón et al., 2014), suggests a relatively important role for genetic drift in driving scenarios of diversification.

Here, we characterize the evolutionary processes underlying the cranial and mandibular diversity across all *Homo*. After quantifying and visualizing variation, we use tests developed from quantitative evolutionary theory to analyze the relative roles of genetic drift and selection within *Homo*, with drift as the null hypothesis. A rejection of drift indicates that morphology is too diverse for divergence to have occurred through random forces alone, thus pointing to a role for adaptation. When present, we then reconstruct the pattern of selection necessary to produce the differences between taxa, identifying the specific morphological regions most likely shaped by selective pressures. These analyses are performed hierarchically to focus on the relationships between temporally successive hominins. We first examine evolutionary process across all of *Homo*. Then, we focus in on the relationships between temporally successive species, at different levels, as well as different geographical populations, to include all possible logical comparisons. In this context, our objectives are to: 1) characterize cranial and mandibular diversity within *Homo* (size and shape), 2) determine whether genetic drift is responsible for this diversity, and 3) explore possible correlations between our results and major evolutionary events, morphological changes and adaptive hypotheses within our genus. Importantly, while this approach cannot predict phylogenetic relationships, it can be used to test hypotheses relating to selective forces acting in human evolution and to investigate causes underlying divergence and ancestor-descendant relationships.

## Materials

Data were collected from the following fossil *Homo* specimens: *Homo sp.* (A.L.666-1, KNM-BC 1, KNM-ER 42703, LD 350-1 [cast], OH 65 [cast], Stw 53 [cast]); *Homo habilis* (KNM-ER 1501, KNM-ER 1802, KNM-ER 1805, KNM-ER 1813, OH 7, OH 13, OH 24, OH 37, OH 62); *Homo rudolfensis* (KNM-ER 1470, KNM-ER 1801, KNM-ER 1482, KNM-ER 3732, KNM-ER 60000 [cast], KNM-ER 62000 [cast], UR 501); *Homo naledi* (DH1, DH2, DH3, DH5); *Homo erectus* (D211 [cast], D2280 [cast], D2282 [cast], D2600 [cast], D2700 [cast], D2735 [cast], KNM-BK 67, KNM-BK 8518, KNM-ER 992, KNM-ER 3733, KNM-ER 3734, KNM-ER 3883, KNM-ER 42700, OH 9, OH 22, Sangiran 1b, Sangiran 4, SK 15, SK 45, SK 847); Middle Pleistocene *Homo* (Bodo [cast], KNM-ER 3884, KNM-ES 11693, LH 18, Ndutu, SAM-PQ-EH 1); Early *Homo sapiens* (Border Cave 1, Border Cave 2, Border Cave 5, Mumbwa, SAM-AP 4692, SAM-AP 6222 KRM1B 41815,Tuinplaas 1). All data, extant and fossil, were collected from NextEngine generated 3D surface scans of the original material or casts in the form of three-dimensional landmarks plotted on the reconstructed 3D surfaces. A total of 33 standard landmarks were extracted from the mandibles and crania (Fig. 1, Table 1). For the comparative sample, landmarks were collected from the left side. For the fossils, all available data were collected and averaged when necessary. These landmarks were used directly in a suite of General Procrustes Analyses. Interlandmark distances were calculated for use in the neutrality tests and multivariate analyses (see Table 2 and Table 3 for lists of fossils utilized in each analysis). Conventional species names and affiliations were used to classify specimens. New specimen affiliations by Spoor and colleagues were considered (Spoor et al., 2015). We use the term Middle Pleistocene *Homo* loosely to denote specimens attributed to the taxa *Homo antecessor, Homo heidelbergensis,* and *Homo rhodesiensis,* as well as archaic *H. sapiens*. We limit our dataset to adult and late juvenile individuals. Due to the fragmentary nature of these specimens, multiple analyses were performed on different regions of the skull, designed to maximize specimen number in some and shared number of variables in others. Landmarks were explicitly chosen based on their repeatability on fossils with varying degrees of preservation. Similarly, specimens were omitted from an analysis if distortion, damage or lack of visible repeatable sutural landmarks were deemed as factors.

**Fig. 1.**
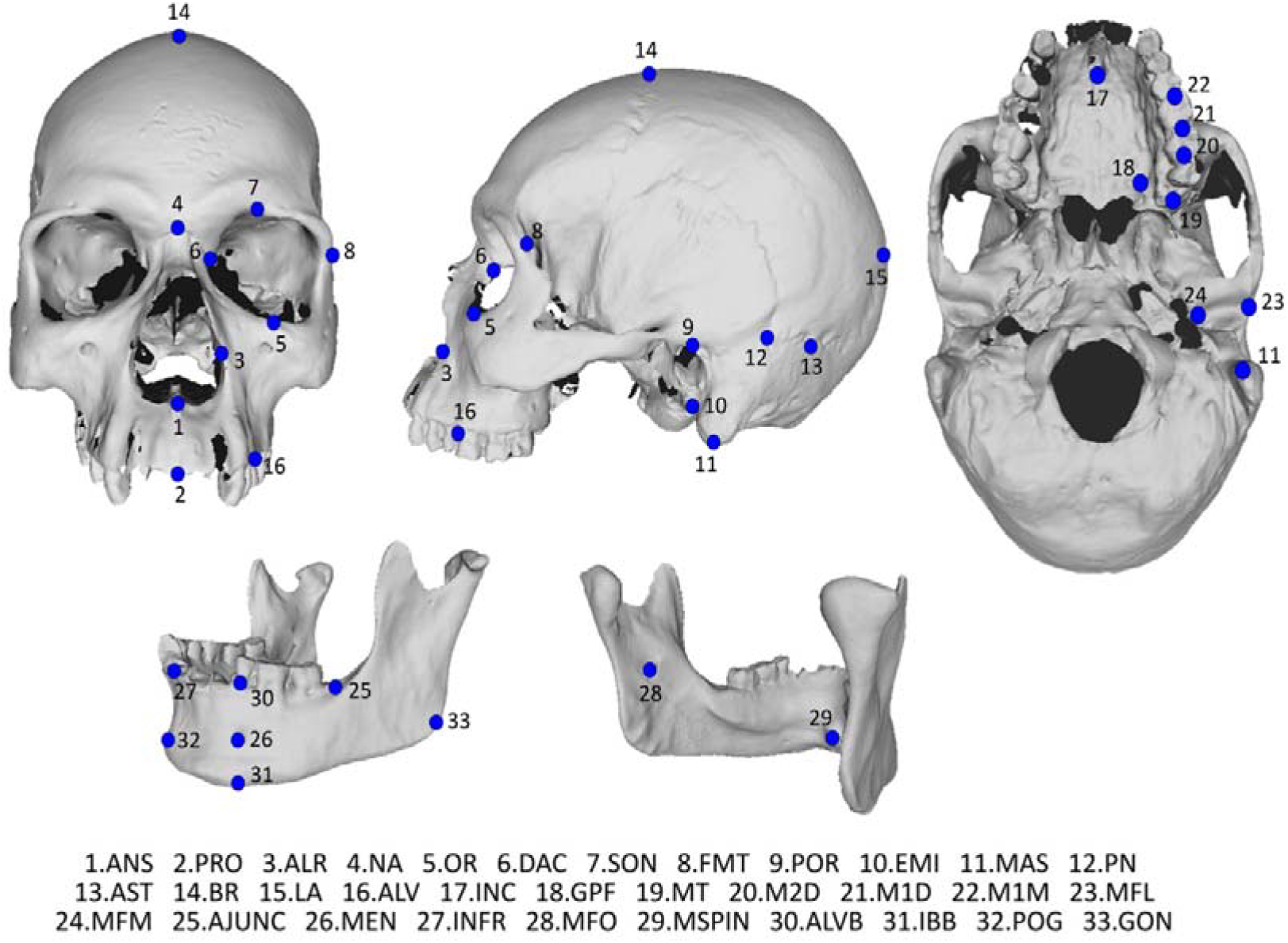
Cranial and mandibular landmarks employed in this study. Landmark definitions and abbreviation descriptions are given in Table 1.

**Table 1.** Standardized landmarks recorded from crania and mandibles. Interlandmark distances are drawn from these landmarks for each analysis ^a^

| Landmark abbreviation | Landmark | Landmark definition |
| --- | --- | --- |
| Cranial landmarks |  |  |
| ANS | Anterior nasal spine | The most anterior midpoint of the anterior nasal spine of the maxilla |
| PRO | Prosthion | The most anterior point in the midline of the maxillary alveolar process |
| ALR | Alare | The most lateral point on the nasal aperture |
| NA | Nasion | The point at the intersection of the nasofrontal suture and the midsagittal plane |
| OR | Orbitale | The most inferior point on the midpoint of the lower edge of the orbit |
| DAC | Dacryon | The point of intersection of the frontolacrimal and lacrimomaxillary sutures |
| SON | Supraorbital notches | The most lateral point on the supraorbital notch |
| FMT | Frontomolare temporale | The most lateral point on the frontozygomatic suture |
| POR | Porion | The most superior point on the margin of the external auditory meatus |
| EMI | External auditory meatus inferior | The most inferior point on the margin of the external auditory meatus |
| MAS | Mastoidale | The most inferolateral point on the mastoid process |
| PN | Parietal notch | The indentation or angle between the squamous and petrous parts of the temporal bone, taken on the superior border of the squama temporalis |
| AST | Asterion | The junction of the lambdoid, parietomastoid and occipitomastoid sutures |
| BR | Bregma | The midline junction of the coronal and sagittal sutures |
| LA | Lambda | The midline junction of the sagittal and lambdoid sutures, taken in the midline |
| ALV | Alveolare | The most anterior point on the alveolus of the M1 (maxillary) |
| INC | Incisivon | The most posteroinferior point on the border of the incisive foramen |
| GPF | Greater palatine foramen | The most posterolateral point on the border of the greater palatine foramen |
| MT | Maxillary tuberosity | The most distal point on the maxillary alveolar process |
| M2D | Distal M2 | The most distal point on M2 |
| M1D | Distal M1 | The most distal point on M1 |
| M1M | Mesial M1 | The most mesial point on M1 |
| MFL | Lateral mandibular fossa | The most lateral point on the mandibular fossa |
| MFM | Medial mandibular fossa | The most medial point on the mandibular fossa |
| Mandibular landmarks |  |  |
| AJUNC | Inferior anterior ramus | The junction of the anterior border of the ramus and alveolus |
| MEN | Mental foramen | The most anteroinferior edge of the mental foramen |
| INFR | Infradentale | The most central point on the mandibular alveolus |
| MFO | Mandibular foramen | The most posteroinferior aspect on the mandibular foramen |
| MSPIN | Superior mental spine | The most superior aspect on the mental spine |
| ALVB | Alveolar border of body | The most superior point on the alveolus directly above the mental foramen |
| IBB | Inferior border of body | The most inferior point on the mandibular corpus directly below the mental foramen |
| POG | Pogonion | The most anterior projection of bone on the mental symphysis |
| GON | Gonion | The junction of the ramus and inferior border of body |

<sup>a</sup> Landmarks and distances are derived from laser surface scans taken by LS. Landmarks are adapted from Ackermann, 1998; Harvati, 2003; Williams and Richtsmeier, 2003; Lockwood et al., 2004; Smith, 2009; von Cramon-Taubadel, 2009; Willmore et al., 2009.

Comparative extant cranial and mandibular material consists of an African *Homo sapiens* sample (N=100) and *Pan troglodytes* sample (N=80) of roughly equal adult males and females. The African *Homo sapiens* sample consists of specimens from the Raymond Dart Collection (RDC) at the University of the Witwatersrand, Johannesburg, and the Iziko Museums of South Africa (SAM), Cape Town, South Africa. The RDC sample (N=50) is a cadaver collection with known sex, age and population group. The individuals from this collection are all identified as Sub-Saharan African. The SAM sample (N=50) is an archaeological collection, with the majority of these individuals categorised as Khoesan, and some associated with dates in the late Holocene. These two collections were chosen for two reasons: 1) the Holocene modern human collection at Iziko museum represents a fairly homogenous population at the extremes of human variation with very limited sexual dimorphism; and 2) the Raymond Dart collection at the University of the Witwatersrand, represents a very diverse mix of African groups, with corresponding variation in body size/shape and dimorphism. Thus, the combination of these two samples provides a satisfactory picture of sub-Saharan African human variation, and is an appropriate model for diversity in the Pleistocene *Homo* sample investigated here.

## Methods

### Evaluating morphological distance between specimens

Mahalanobis’ generalised distance statistic (*D*^2^), a multivariate approach, was used to explore the morphological distances between fossil *Homo* specimens to assess the variability within the sample, substituting a variance/covariance (V/CV) matrix of *H. sapiens* as an estimate of within species variation (as per Ackermann, 2003). Mahalanobis’ Distance values between specimens were calculated from interlandmark distances, scaled to the geometric mean, using MATHEMATICA^™^ v8. Shared variables between fossil specimens and extant samples are converted into vector form. These vectors are used in the following equation to calculate Mahalanobis’ distances: *D*^2^ *=* (*x*_1_ – *x*_2_) *′ V*^-1^ (*x*_1_ – *x*_2_), where *D*^2^ is the Mahalanobis’ distance between specimens one and two, *x*_1_ is the vector of values for specimen one, *x*_2_ is the vector of values for specimen two, and *V*^-1^ is the inverse of the variance/covariance matrix of the extant *Homo sapiens* model population. A series of Principal Coordinates (PCoord) Analyses was then performed on the matrices of Mahalanobis’ distances for each analysis to visualize the morphological differences and overall variation among the fossil individuals. All PCoord analyses were performed in PAST v2.17 (Hammer et al., 2001). To evaluate significance, frequency distributions of expected Mahalanobis’ distances were calculated for the *Homo sapiens* model population using its own V/CV matrix, as well as a *Pan troglodytes* V/CV matrix (Ackermann, 2003). Fossil distances are considered significantly different when they exceed the 95^th^ percentile of values for the generated frequency distributions. This method allows us to measure how well intra-specific variation is evaluated in extant species using a covariance matrix calculated from its own species or a closely-related species.

### Investigating the effect of size

Generalized Procrustes Analysis (GPA), a superimposition method, was performed on three-dimensional landmark data (Fig. 1; Table 1) to investigate the effect of size and size-related shape on *Homo* crania and mandibles (Gower, 1975; Rohlf and Slice, 1990; Rohlf, 1999). Landmark configurations are standardised, translated and scaled, to the same centroid size, and rotated so that the summed squared distances between the landmarks and their corresponding sample mean is minimized. The resultant transformed coordinates of these superimposed landmarks are the Procrustes shape coordinates, represented as points in Kendall’s shape space, containing information about the shape of the original landmark configurations. To visualize the shape differences between specimens and to identify the major axes (patterns) of variance, principal components analysis (PCA) was performed on the covariance matrix of the transformed Procrustes shape coordinates. Shape variability along each principal component was further evaluated using reconstructed wireframes (not presented) based on the original landmark configurations and the resultant shape change was then assessed. Allometry, the correlation of size and shape, was assessed using pooled within-group multivariate regressions of shape (Klingenberg, 1996, 1998, 2016; Monteiro, 1999; Mitteroecker et al., 2003). This was done by regressing the Procrustes coordinates on log centroid size. The significance of this potential correlation was assessed using a permutation test (10,000 runs) against the null hypothesis of independence between the dependent and independent variables. All analyses were performed using the geometric morphometrics software MorphoJ version 1.05f (Klingenberg, 2011).

### Testing the null hypothesis of genetic drift

According to the quantitative genetic theory of Lande (Lande, 1997, 1979, 1980), the neutral model of evolution is shown by the equation: *E(B*_*t*_*) = G(t / N*_*e*_), where *E(B*_*t*_) is the expected between population variance/covariance (V/CV) matrix, *t* is the number of generations, *G* is the additive genetic V/CV matrix and *N*_*e*_ is the population size. We use *E(B*_*t*_) as an expectation operator to emphasize that the scenario whereby *B*_*t*_ is exactly equal to *G(t / N*_*e*_) is highly unlikely. Quantitative theory has shown that the phenotypic within-group V/CV matrix (*W*) is correlatively similar to *G*, thus allowing us to substitute *W* for *G* (Cheverud, 1988). Therefore, if random genetic drift has shaped the diversity seen within *Homo*, a proportional relationship should exist between the patterns of *Homo* between-group variation and the within-group extant *Homo sapiens* variation (*B* ∝ *W*). To assess this relationship, we regress the logged between-group eigenvalues (*B*), calculated as the variance among group mean differences between fossil populations, onto logged within-group eigenvalues (*W*), obtained from principal components calculated from the extant covariance matrices substituted as models for within-population variability. If populations have diversified through random genetic drift then the regression slope will not be distinguishable from a slope of 1.0 (at a 0.05 significance level), indicating that the pattern of variance within and between these groups is comparable and changes in magnitude are mostly due to scaling. A non-proportional relationship or rejection of drift indicates that morphology is too variable for divergence to have occurred through random forces alone and non-random forces, such as directional selection, are likely to be at work. Rate tests performed in a previous study using a subset of these data support the capacity of the slope test to distinguish between random genetic drift and selection (Schroeder et al., 2014). It is important to note that these slope tests are also able to detect the difference between random selection and random genetic drift. This statement holds true unless random selection acts in a manner that distributes it exactly along the lines of the within group covariation, i.e. that selection is exactly proportional to the covariation in the population – which is unlikely. All analyses were performed in R version 3.0.1.

### Reconstructing selection

When a null hypothesis of genetic drift is rejected, we reconstruct the selection necessary to produce the differences in observed population means. The methodological approach derives from the quantitative evolutionary theory of Lande (Lande and Arnold, 1983) and is determined by the following relationship: *β = W*^-1^[ *z*_*i*_ *− z*_*j*_], where *β* is the differential selection gradient/vector summed over generations, *W*^-1^ is the inverse of the within-species phenotypic V/CV matrix (again used here as a proxy for the additive genetic covariance matrix), and [*z*_*i*_ *− z*_*j*_] is the difference in means between species *i* and *j*, in this case the fossil species being compared. As before, we use the V/CV matrices from an extant *Homo sapiens* sample substituted as a model for fossil within-species variation. The reconstructed selection vectors are used to investigate the direction or pattern of selection, (less so the magnitude of selection), acting to differentiate *Homo* groups. The direction of selection, positive or negative, is subject to our expectation of the basic ancestor-descendent relationships among these groups. The magnitude of selection is strongly dependent on the estimated covariance matrix structure and therefore we interpret these results with caution. We highlight strongly negative (<-1) and strongly positive (>1) gradients, however these levels are not statistically evaluated. Schroeder and colleagues (2014) performed matrix corrections to account for the error in estimated covariance matrices and investigate the possible impact that this error may have in the calculation of selection gradients. Although these corrections affected the magnitude of selection, the resultant effect on the pattern/direction of selection was found to be negligible (Schroeder et al., 2014).

## Results

Surface scans and landmarks were collected from 48 original fossil specimens of *Homo* and 12 high quality casts (see *Materials and Methods*). Multiple analyses were performed on both linear measurements and three-dimensional landmark data (visualization of landmarks in Fig. 1; landmark descriptions in Table 1). These analyses were designed to maximize specimen number and/or shared variables in order to analyze as much cranial and mandibular material as possible. Multivariate analyses (Mahalanobis’ distances) and tests for genetic drift were applied separately to fourteen different sets of interlandmark distances (10 cranial, 4 mandibular), representing all regions of the cranium and mandible (described in Table 2). Geometric morphometric analyses were performed on eleven subsets of landmarks (7 cranial and 4 mandibular; Table 3). For all analyses, specimen choice was dependent on the availability of landmarks. Some specimens and variables were omitted from analyses due to the lack of visible landmarks, preservation or distortion. An extant *Homo sapiens* sample (N=100) was used as a comparative taxon in the geometric morphometric analyses, and as a best-fit model of intra-specific variability both for calculation of Mahalanobis’ distances and in the tests for genetic drift, under the assumption that each extinct taxon had similar within-species covariance structures as *Homo sapiens*. Chimpanzees (*Pan troglodytes*) are not seen to be an appropriate model of covariance for evaluating *Homo*; regardless, tests for genetic drift using a chimpanzee model provided comparable results to those using *Homo sapiens* in an earlier study on a reduced data set (Schroeder et al., 2014). Mahalanobis’ distances were calculated on interlandmark distances scaled to the geometric mean. Neutrality tests were performed on unscaled data to evaluate both size and shape change.

**Table 2.**
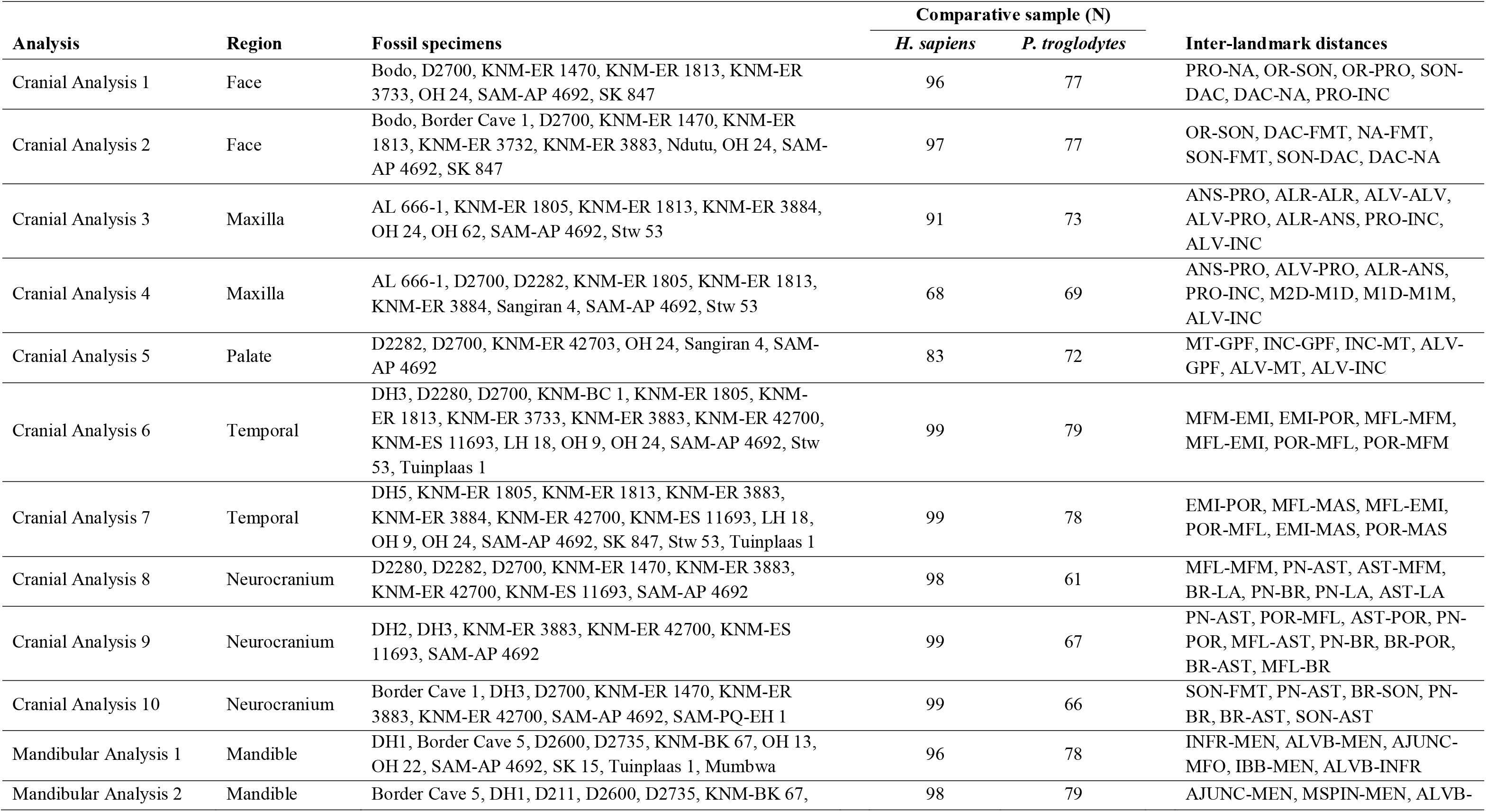
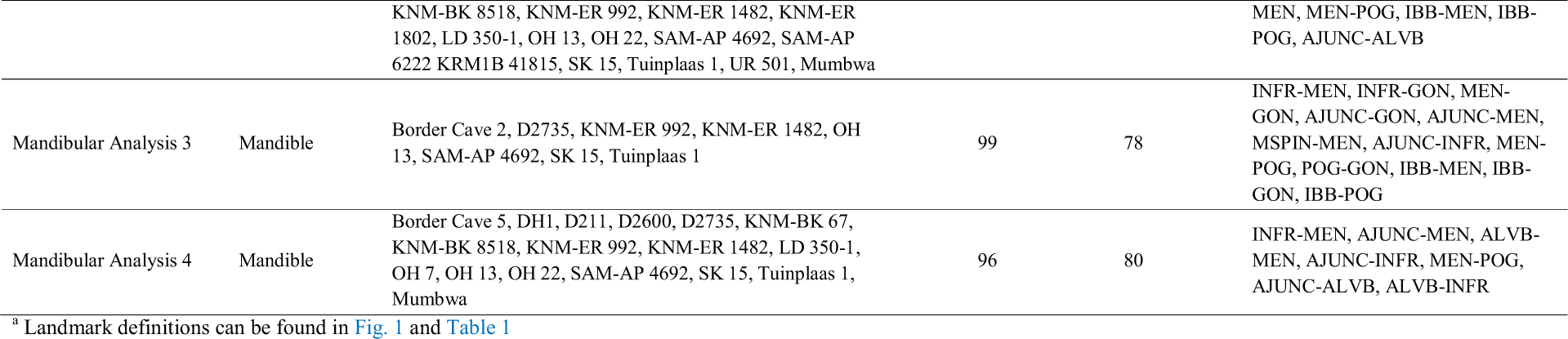
Description of each multivariate analysis (for multivariate and neutrality tests) including list of fossil specimens, number of comparative individuals and inter-landmark distances ^a^

### Multivariate assessment of variability

Mahalanobis’ distance values and 95^th^ percentile significance values for each analysis are presented in SOM Dataset 1. Matrices of Mahalanobis’ distances were used in a series of Principal coordinates (PCoord) analyses to visualize the morphological differences between fossil specimens (Fig. 2 and SOM Fig. S1). The PCoord analyses of the Mahalanobis’ distances (*D^2^*) calculated between fossil specimens on different subsets of cranial and mandibular traits depict relatively clear species clusters and illustrate the variability among Pleistocene *Homo* specimens, especially within *H. erectus*.

**Fig. 2.**
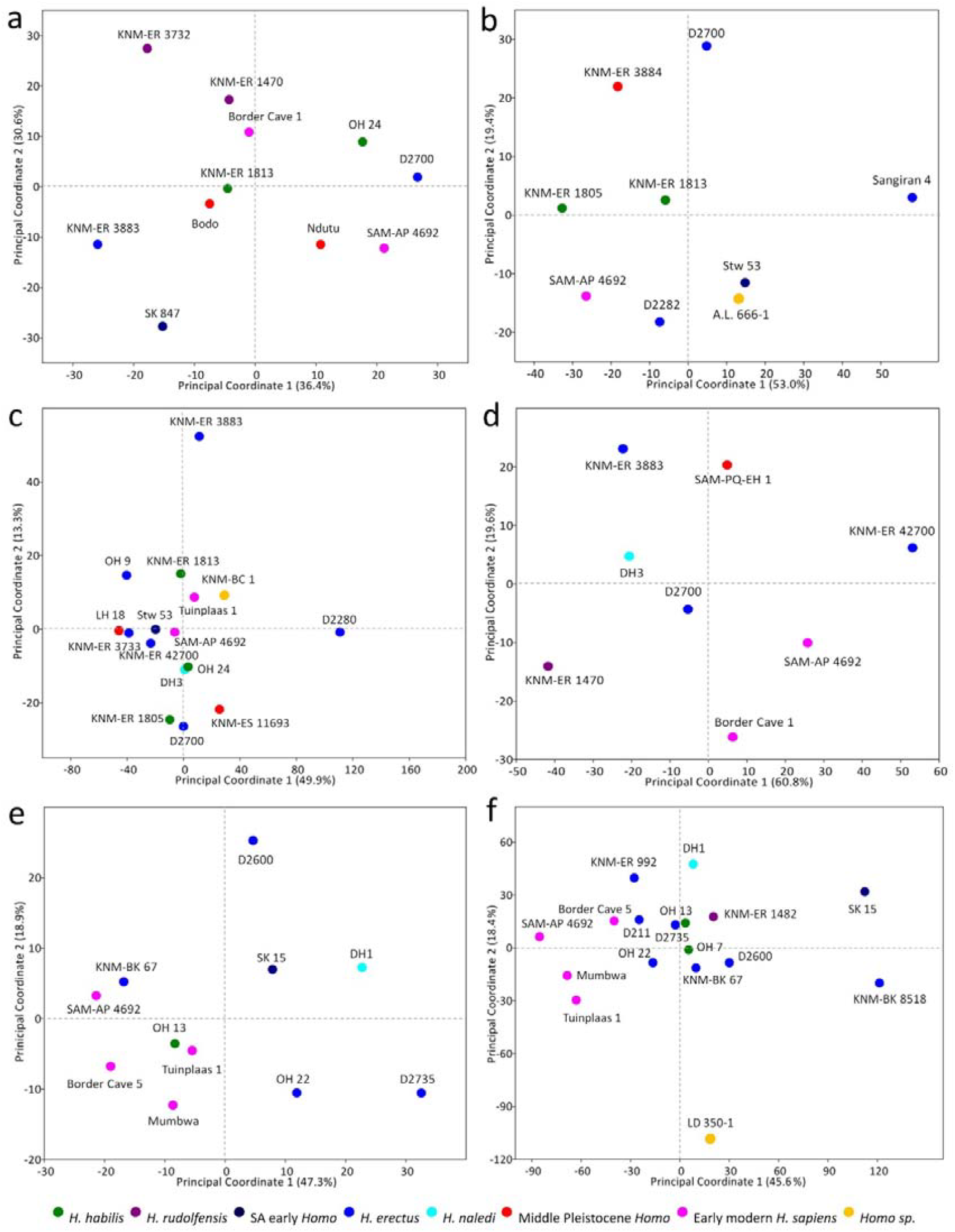
A subset of principal coordinates plots of Mahalanobis’ distances between fossil specimens, using a *Homo sapiens* variance/covariance model. The remaining principal coordinates plots are illustrated in SOM Fig. S1. Analyses were performed on scaled interlandmark distances (See Table 2 for further information). Percentage of variance explained by each principal coordinate is displayed on each plot. Matrices of Mahalanobis’ distances (*D*^*2*^) can be found in SOM Dataset 1. (a) Cranial Analysis 2 (face). Large, significantly different, *D*^*2*^ values are identified between KNM-ER 3732 and SK 847, KNM-ER 3732 and SAM-AP 4692, D2700 and KNM-ER 3883, D2700 and SK 847, as well as SK 847 and OH 24. (b) Cranial Analysis 4 (maxilla). Sangiran 4 is significantly different from SAM-AP 4692, KNM-ER 3884, KNM-ER 1805, D2282, and D2700. A.L.666-1 is closely associated with Stw 53. (c) Cranial Analysis 6 (temporal). D2280 is significantly different from all other specimens, notably LH 18 and OH 9. KNM-ER 3883 is also an outlier, however, this specimen still shows some affinities with KNM-ER 1813 and KNM-BC 1. (d) Cranial Analysis 10 (neurocranium). Large significantly different *D*^*2*^ values are found between KNM-ER 1470 and KNM-ER 42700, DH3 and KNM-ER 42700, as well as KNM-ER 3883 and KNM-ER 42700. (e) Mandibular Analysis 1. D2600 is significantly different from all other specimens. Other notable significantly different values are between D2735 and SAM-AP 4692, and D2735 and KNM-BK 67. DH1 is closely associated with SK 15 and OH 22. (f) Mandibular Analysis 4. LD 350-1 is significantly different from all specimens, with the exception of Tuinplaas 1 and KNM-BK 8518. Other significant differences can be found between SK 15, KNM-BK 8518 and early modern *H. sapiens.* The plot shows an overlap between *H. erectus, H. habilis* and *H. rudolfensis.*

#### Cranial analyses

For analyses of the face (cranial analyses 1, 2), *Homo rudolfensis* specimens KNM-ER 1470 and KNM-ER 3732, and Dmanisi *Homo erectus* specimen D2700 are consistently significantly different from SK 847, KNM-ER 3733, and KNM-ER 3883 (Fig. 2a; SOM Fig. S1A). For analyses of the maxilla (cranial analyses 3, 4, 5), specimens A.L.666-1, Sangiran 4 and KNM-ER 42703 are shown to be significantly different from other specimens (Fig. 2b; SOM Fig. S1B-C). For analyses of the temporal (cranial analyses 6, 7), *H. erectus* specimens D2280, KNM-ER 3883 and OH 9 are significantly different from most other specimens (Fig. 2c; SOM Fig. S1D). In neurocranial analyses 8 and 9, large *D*^*2*^ values are recorded between *H. erectus* specimens (D2800, D2282, KNM-ER 3883 and KNM-ER 42700) and all other specimens (SOM Figs S1E-F). Neurocranial analysis 10 also produces large *D*^*2*^ values between KNM-ER 42700 and all other specimens, with the exception of SAM-AP 4692 (Fig. 2d). KNM-ER 1470 is significantly different from KNM-ER 42700, SAM-AP 4692 and SAM-PQ-EH 1.

#### Mandibular analyses

For mandibular analysis 1, D2600, *Homo naledi* specimen DH1, D2735, OH 22 and SK 15 are significantly different from all other specimens (Fig. 2e). Mandibular analyses 2 and 4 depict a similar pattern where LD 350-1, KNM-BK 8518 and D2600 are consistently significantly different from most other specimens (Fig. 2f; SOM Fig. S1G). In mandibular analysis 3, D2735 and KNM-ER 1482 are shown to be significantly different from other specimens (SOM Fig. S1H).

### Geometric morphometrics

A summary of the results for all eleven analyses (four mandibular [GPA 1-4] and seven cranial [GPA 5-11]) can be found in Table 3. Fig. 3 and SOM Fig. S2 display the principal component plots (of principal component one and two for each analysis). The first two components explain between 43% and 79% of the shape variation among specimens for these analyses. The shape changes associated with each principal component are described in Table 3. The first principal component (PC1) is the most taxonomically diagnostic in all analyses. A *Homo sapiens* sample was included in the analysis to provide context. We also include a sample of *Pan troglodytes* as an outgroup for a subset of analyses to explore the effect that this could have on the interpretation of our results (SOM Fig. S3).

**Fig. 3.**
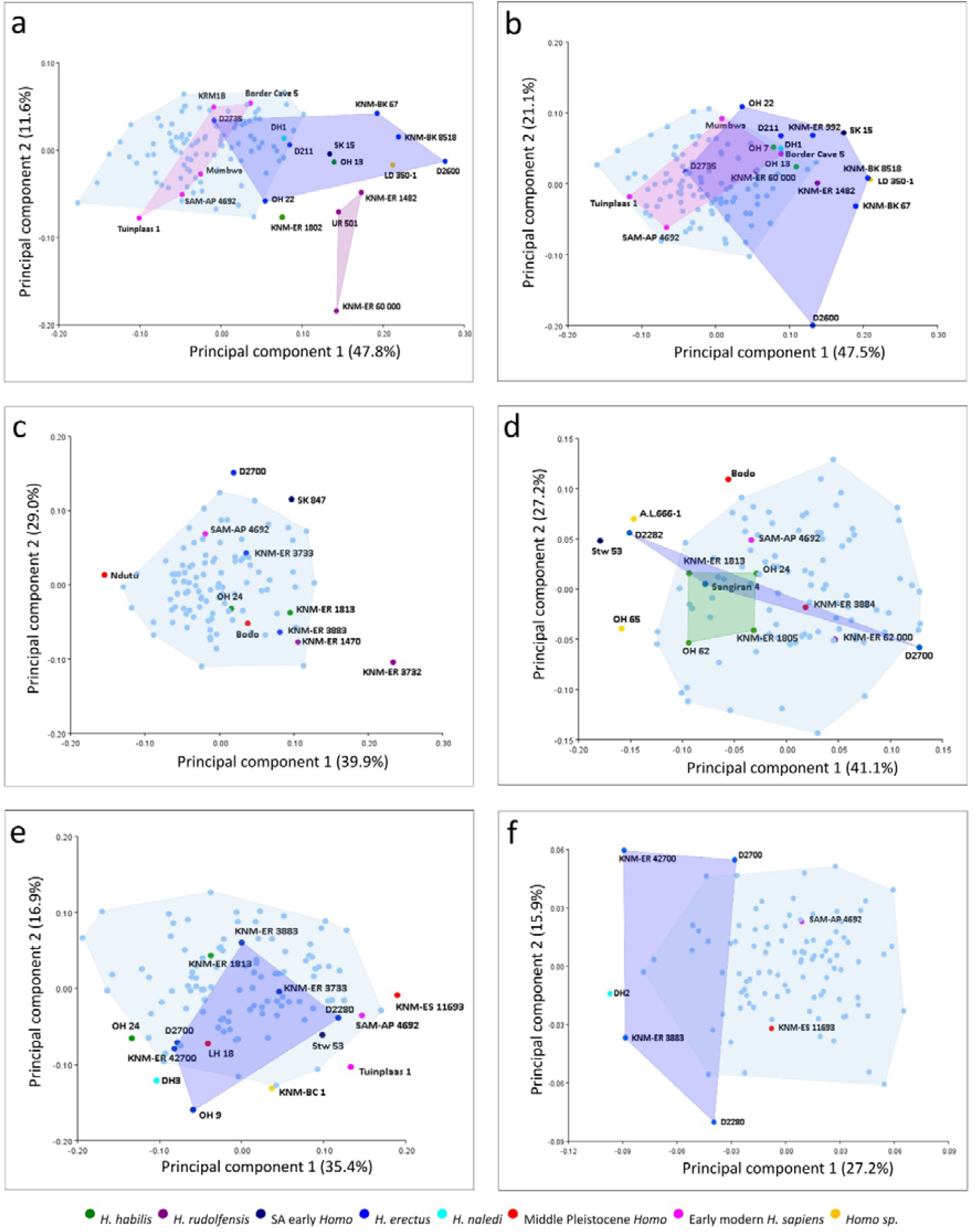
Principal component plots of PC1 and PC2 for a subset of Generalized Procrustes analyses (GPA). The remaining principal components plots are illustrated in SOM Fig. S2. A summary of all GPA results is given in Table 3. The percentage of variance explained by each principal component is displayed on each plot. (a) GPA 1 – mandible. Species convex hulls are separated along PC1. Most Pleistocene *Homo* specimens fall within the *H. erectus* convex hull, with the exception of *H. rudolfensis* specimens and KNM-ER 1802. KNM-ER 60000 is an outlier along PC2. (b) GPA 2 – mandible. There is a fair amount of overlap between species convex hulls. All Pleistocene *Homo* specimens are contained within the convex hull of *H. erectus*, with the exception of LD 350-1 which falls just outside of the range. D2600 is an outlier along PC2. (c) GPA 5 – upper face. Most specimens fall within the *H. sapiens* range, except for Ndutu, SK 847, D2700 and KNM-ER 3732. (d) GPA 6 – maxilla. Dmanisi *H. erectus* shows the most variability along PC1. A.L.666-1 is closely associated with D2282 and Stw 53 in shape space, and OH 65 along PC1. The *H. habilis* convex hull is enclosed within the *H. sapiens* range. (e) GPA 8 – temporal. Most specimens are contained within the *H. sapiens* convex hull, with the exception of OH 24, DH3, OH 9, KNM-BC 1, Tuinplaas 1 and KNM-ES 11693. (f) GPA 11 – neurocranium. *H. erectus* is most variable along PC2. DH2 falls just outside the convex hull of *H. erectus* along PC 1.

**Table 3.**
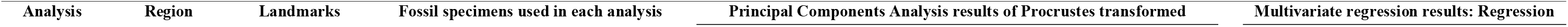
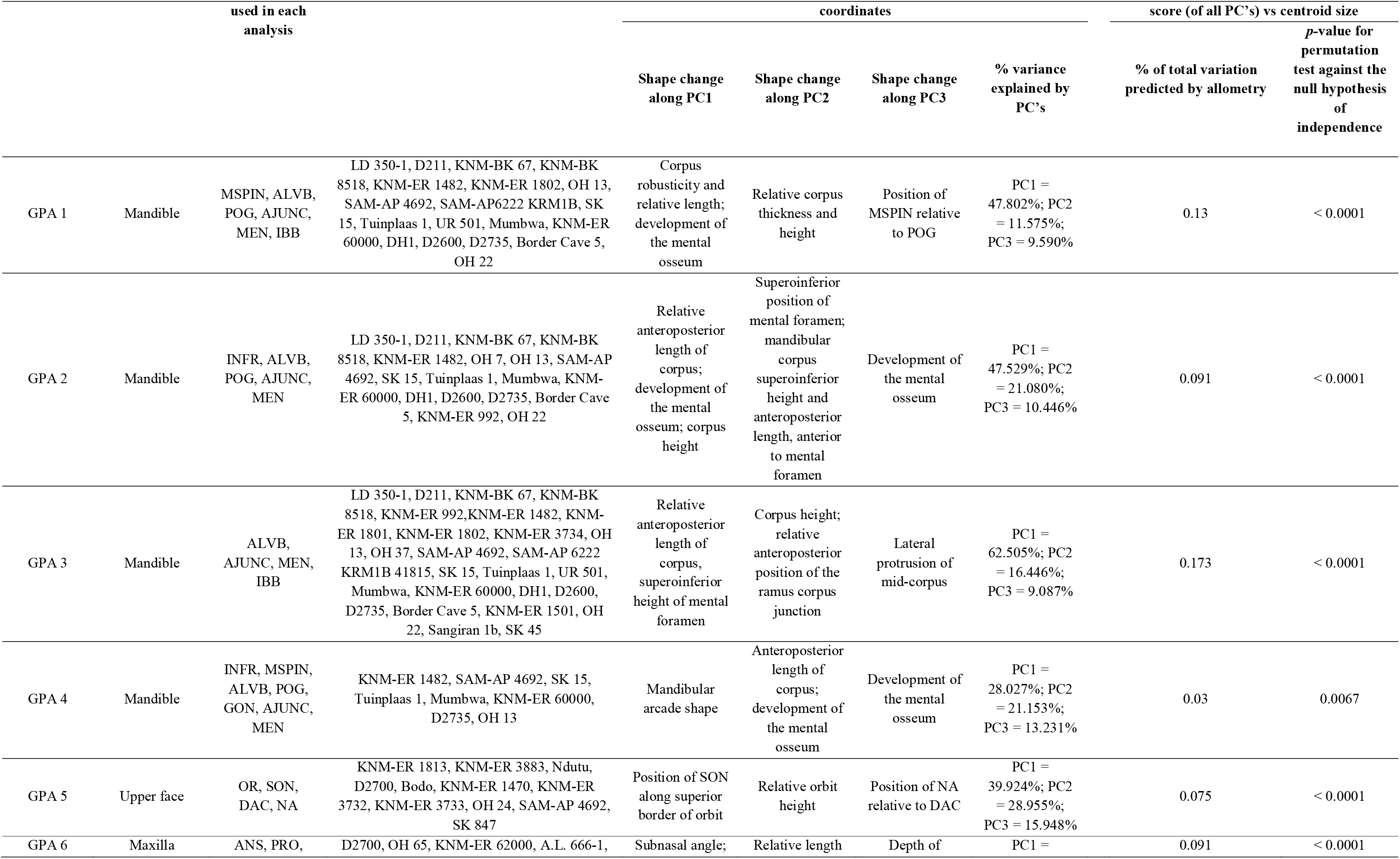
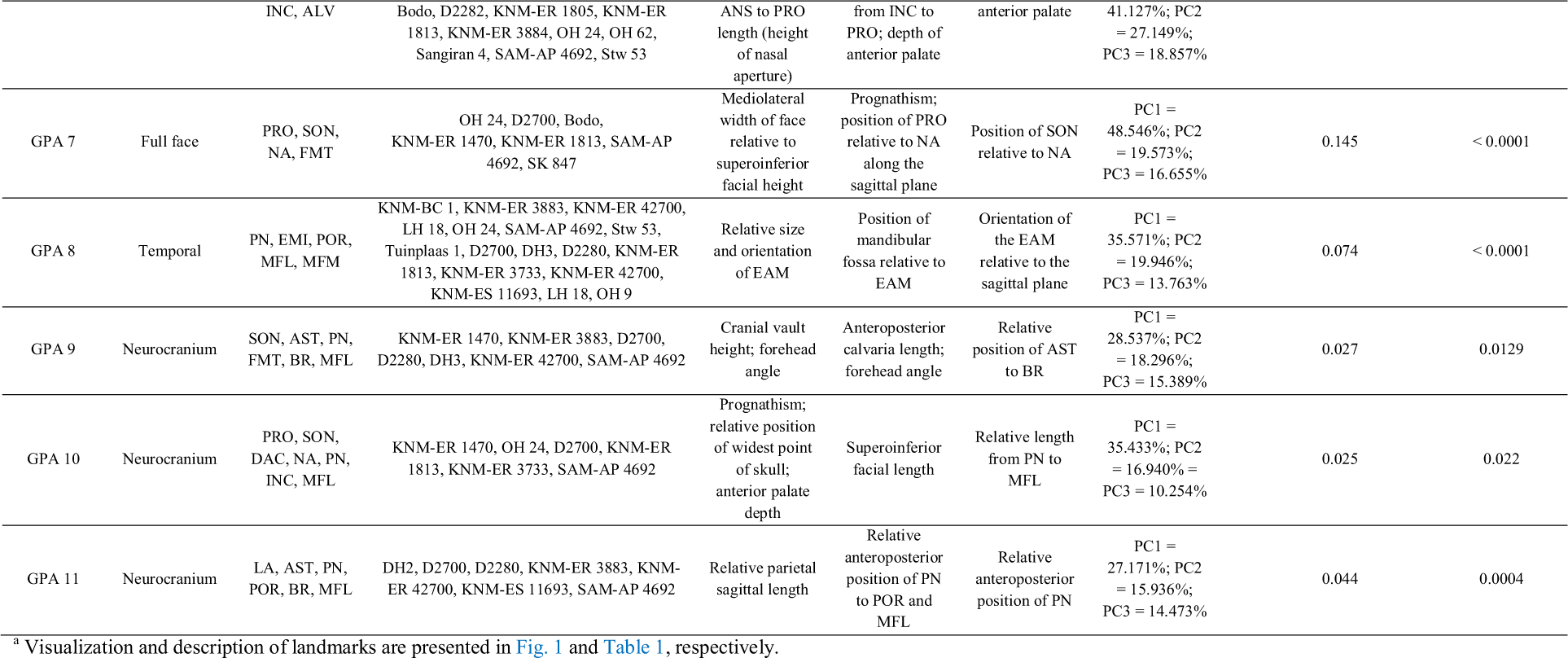
Geometric morphometric results summary and analysis description^a^

#### Mandibular analyses

In general, species convex hulls separate along PC1 in all mandibular analyses, however significant overlap is observed. In GPA 1, most fossil *Homo* specimens are contained within the *H. erectus* convex hull, once again highlighting the diversity of this hypodigm (Fig. 3a). Interestingly, all *H. rudolfensis* specimens are excluded from this shape space, as well as KNM-ER 1802, a specimen traditionally placed within *H. rudolfensis* and recently reclassified as *H. habilis* (Antón et al., 2014; Spoor et al., 2015). These specimens, especially KNM-ER 60000, are separated from all others along PC2, which reflects relative corpus thickness and height. PC1 reflects corpus robusticity, relative corpus length and the development of the mental osseum. For GPA 2, the amount of overlap is substantial, with the vast majority of fossil *Homo* specimens falling within the *H. erectus* range (Fig. 3b). LD 350-1 falls just outside of this range. Outlier D2600 is separated from other specimens along PC2, which reflects a change in relative corpus height, length and mental foramen position. PC1 also corresponds to a change in relative corpus height and length, as well as development of the mental osseum. GPA 3 depicts a similar pattern to GPA 2, with most fossil *Homo* specimens falling within the convex hulls of *H. erectus* and *H. sapiens,* as well as a fair amount of species overlap (SOM Fig. S2A). GPA 4, D2735 falls within the *H. sapiens* convex hull, with all other specimens falling outside, separated along PC2 (SOM Fig. S2B).

#### Cranial analyses

In cranial analyses of the face (GPA 5, 6, 7), most specimens fall within the *H. sapiens* convex hull. The exceptions are as follows: in GPA 5, KNM-ER 3732 and Middle Pleistocene *Homo* specimen Ndutu are separated from all other specimens along PC1, D2700 and SK 847 are separated from the others along PC2 (Fig. 3c); in GPA 6, Dmanisi *H. erectus* shows the most variability along PC1, with both specimens falling outside the convex hull of *H. sapiens* (Fig. 3d). A.L.666-1, Stw 53 and OH 65 separate from all other specimens along PC1 (Fig. 3d); in GPA 7, OH 24 and KNM-ER 1470 are not contained in the *H. sapiens* convex hull, with KNM-ER 1470 and OH 24 falling at the positive extreme of PC1 and PC2, respectively (SOM Fig. S2C). In the analysis of the temporal bone (GPA 8), DH3, OH 24, KNM-BC 1, KNM-ES 11693 and Tuinplaas 1 all fall outside of the *H. sapiens* range (Fig. 3e). This plot is not particularly taxonomically diagnostic. The focus of the final three cranial analyses is neurocranium shape (GPA 9, 10, 11). There is moderate species overlap, with species separating along PC1, which reflects relative vault height, length and breadth (Fig. 3f; SOM Figs S2D-E). For GPA 9, KNM-ER 42700 is an outlier at the positive extreme of PC2, reflecting relative vault length and forehead slope (SOM Fig. S2.D). For GPA 11, DH2 falls outside of the convex hull of *H. erectus,* along PC1, which corresponds to relative parietal sagittal length (Fig. 3f).

### Testing the null hypothesis of genetic drift

Following the quantitative evolutionary theory of Lande (Lande, 1977, 1979, 1980) and the methodological approach of Ackermann and Cheverud 2004 and Schroeder et al. 2014, the null hypothesis of genetic drift is tested, i.e. the hypothesis that between-group and within-group phenotypic variation should be proportional under a neutrally evolving model. Regression results of logged between-group to logged within-group variation to test the deviation from a slope of 1.0 are given in SOM Table S1 and summarized in Table 4. The results indicate that for 95% of all analyses, performed across all taxa and skull regions, the null hypothesis of genetic drift cannot be rejected. This is particularly apparent in analyses of the neurocranium, where all 39 comparisons are consistent with random genetic drift. This suggests that differences in the pattern of covariance among neurocranial traits are negligible, regardless of which taxa are being compared. However, it is important to note here that a failure to reject drift does not completely remove the possibility that non-random processes were acting, but rather indicates that any effect of these processes cannot be distinguished from divergence due to drift. Furthermore, the structure of the test makes it difficult to reject drift when few traits are being compared, because the number of measurements (number of PCs) is directly related to the degrees of freedom. The power of the test is also influenced by the strength of the correlation between two taxa diverging under a model of neutrality, with the strength of the correlation decreasing the longer the split time between taxa. For these reasons, any significant deviation from a slope of 1.0 will likely signify selection. On the other hand, it is possible that given the large number of tests performed and the possibility of Type II errors, a rejection of genetic drift may be a reflection of false positives in the data at a 0.05 significance level. However, we still regard this as a conservative estimate given the lack of power of this test. Because of these issues, it may be prudent to focus on those analyses which have the highest number of traits (cranial analysis 9 and mandibular analysis 3) and therefore a relatively high statistical power (calculated using the pwr.f2.test in the “pwr” package in R v3.2.2 [Champely, 2016]), as well as comparisons with high R^2^ values, indicating a good fit to the model. When this is done, we still cannot reject drift for 51% of all comparisons, supporting our conclusions above.

**Table 4.** Summary results of between-group variance regressed on within-group variance as a test for genetic drift.

| Analysis | Consistent with drift | Rejection of drift |
| --- | --- | --- |

|  | Groups compared | Number of<br>comparisons<br>consistent with drift | Comparison | Slope | R<sup>2</sup> | P-value |
| --- | --- | --- | --- | --- | --- | --- |
| <u>Cranial Analysis 1</u><br>Face | *H. habilis*, *H. rudolfensis*, SA early *Homo*, African *H. erectus*, Dmanisi *H. erectus*, *H. erectus (sensu lato)*, Middle Pleistocene *Homo*, early modern *H. sapiens* | 18/19 | African *H. erectus* - Middle Pleistocene *Homo* | 0.22 | 0.19 | 0.03 |
| <u>Cranial Analysis 2</u><br>Face | *H. habilis*, *H. rudolfensis*, SA early *Homo*, African *H. erectus*, Dmanisi *H. erectus*, *H. erectus (sensu lato)*, Middle Pleistocene *Homo*, early modern *H. sapiens* | 18/19 | *H. habilis* - Dmanisi *H. erectus* | 0.14 | 0.08 | 0.02 |
| <u>Cranial Analysis 3</u><br>Maxilla | A.L.666-1, *H. habilis*, SA early *Homo*, Middle Pleistocene *Homo*, early modern *H. sapiens* | 5/5 | None | — | — | — |
| <u>Cranial Analysis 4</u><br>Maxilla | A.L.666-1, *H. habilis*, *H. rudolfensis*, SA early *Homo*, Dmanisi *H. erectus*, Asian *H. erectus*, *H. erectus (sensu lato)*, Middle Pleistocene *Homo*, early modern *H. sapiens* | 19/21 | A.L.666-1 - Dmanisi *H. erectus*<br>*H. habilis* - Dmanisi *H. erectus* | 1.87<br>0.05 | 0.87<br>0.01 | 0.04<br>0.02 |
| <u>Cranial Analysis 5</u><br>Palate | *H. habilis*, KNM-ER 42703, Dmanisi *H. erectus*, Asian *H. erectus*, *H. erectus (sensu lato)*, early modern *H. sapiens* | 12/12 | None | — | — | — |
| <u>Cranial Analysis 6</u><br>Temporal | KNM-BC 1, *H. habilis*, SA early *Homo*, African *H. erectus*, *H. naledi*, Dmanisi *H. erectus*, Asian *H. erectus*, *H. erectus (sensu lato)*, Middle Pleistocene *Homo*, early modern *H. sapiens* | 28/29 | *H. habilis* - African *H. erectus* | 0.10 | 0.02 | 0.05 |
| <u>Cranial Analysis 7</u><br>Temporal | *H. habilis*, SA early *Homo*, African *H. erectus*, *H. naledi*, Middle Pleistocene *Homo*, early modern *H. sapiens* | 18/20 | Stw 53 - east African *H. erectus*<br>African *H. erectus* - Middle Pleistocene *Homo* | 0.06<br>-0.67 | 0.02<br>0.41 | 0.01<br>0.01 |
| <u>Cranial Analysis 8</u><br>Neurocranium | *H. rudolfensis*, African *H. erectus*, Dmanisi *H. erectus*, *H. erectus (sensu lato)*, Middle Pleistocene *Homo*, early modern *H. sapiens* | 13/13 | None | — | — | — |
| <u>Cranial Analysis 9</u><br>Neurocranium | African *H. erectus*, *H. naledi*, Middle Pleistocene *Homo*, early modern *H. sapiens* | 8/8 | None | — | — | — |
| <u>Cranial Analysis 10</u><br>Neurocranium | *H. rudolfensis*, African *H. erectus*, *H. naledi*, Dmanisi *H. erectus*, *H. erectus (sensu lato)*, Middle Pleistocene *Homo*, early modern *H. sapiens* | 18/18 | None | — | — | — |
| <u>Mandibular Analysis 1</u> | *H. habilis*, SA early *Homo*, African *H. erectus*, *H. naledi*, Dmanisi *H. erectus*, *H. erectus (sensu lato)*, early modern *H. sapiens* | 18/20 | *H. habilis* - Dmanisi *H. erectus*<br>*H. habilis* - *H. erectus (sensu lato)* | 4.17<br>2.21 | 0.91<br>0.94 | 0.02<br>0.03 |
| <u>Mandibular Analysis 2</u> | LD 350-1, *H. habilis*, *H. rudolfensis*, SA early *Homo*, African *H. erectus*, *H. naledi*, Dmanisi *H. erectus*, *H. erectus (sensu lato)*, early modern *H. sapiens* | 38/39 | *H. rudolfensis* - Dmanisi *H. erectus* | -0.42 | 0.11 | 0.04 |
| <u>Mandibular Analysis 3</u> | *H. habilis*, *H. rudolfensis*, SA early *Homo*, African *H. erectus*, *H. naledi*, Dmanisi *H. erectus*, *H. erectus (sensu lato)*, early modern *H. sapiens* | 16/16 | None | — | — | — |
| <u>Mandibular Analysis 4</u> | LD 350-1, *H. habilis*, *H. rudolfensis*, SA early *Homo*, African *H. erectus*, *H. naledi*, Dmanisi *H. erectus*, *H. erectus (sensu lato)*, early modern *H. sapiens* | 27/31 | LD 350-1 - *H. rudolfensis*<br>*H. habilis* - *H. rudolfensis*<br>*H. rudolfensis* - African *H. erectus*<br>*H. rudolfensis* - *H. erectus (sensu lato)* | 0.45<br>0.34<br>0.40<br>0.28 | 0.47<br>0.33<br>0.56<br>0.18 | 0.05<br>0.03<br>0.01<br>0.05 |

In the remaining (5%) cases drift is rejected at a 0.05 significance level (SOM Fig. S4). For cranial analysis 1 (face), a rejection of drift is detected between Middle Pleistocene *Homo* and African *H. erectus.* For cranial analysis 2 (face), drift is rejected between *H. habilis* and Dmanisi *H. erectus.* The slopes for both these analyses are <1.0, and examination of the regression plots indicates that the first one or more PCs show less than expected between-group (fossil) variation, relative to minor PCs (SOM Figs S4A-B). For cranial analysis 4 (maxilla), a rejection of drift is detected between early *Homo* specimen A.L.666-1 and Dmanisi *H. erectus,* as well as *H. habilis* and Dmanisi *H. erectus*. The slope for the comparison of A.L.666-1 and Dmanisi *H. erectus* is >1.0, which appears to be driven by more between-group variation in the first few PCs than expected and less in the minor PCs (SOM Fig. S4C). Conversely, the slope for the comparison of *H. habilis* and Dmanisi *H. erectus* is <1.0, the result of less between-group variation in the first few PCs and more in lesser PCs (SOM Fig. S4D). For cranial analysis 6 and 7 (temporal), drift is rejected between *H. habilis* and African *H. erectus,* South African early *Homo* specimen Stw 53 and *east African H. erectus,* as well as African *H. erectus* and Middle Pleistocene *Homo.* For all analyses, the slopes are <1.0, primarily due to less than expected between-group variation in the first few PCs (SOM Figs S4E-G). In mandibular analysis 1, drift is rejected between *H. habilis* and Dmanisi *H. erectus,* and *H. habilis* and *H. erectus (sensu lato),* with more between-group variation in the first few PCs and less in the minor PCs than expected (SOM Figs S4H-I). For mandibular analysis 2, a rejection of drift is detected between *H. rudolfensis* and Dmanisi *H. erectus*, with less between-group variation in the first few PCs than expected (SOM Fig. S4J). Deviations from genetic drift are detected among four comparisons in mandibular analysis 4. These comparisons are LD 350-1 and *H. rudolfensis, H. habilis* and *H. rudolfensis, H. rudolfensis* and African *H. erectus,* as well as *H. rudolfensis* and *H. erectus (sensu lato).* All slopes are <1.0, showing less than expected between-group variation in the first few PCs (SOM Figs S4K-N).

### Reconstructing patterns of selection

For the fourteen comparisons where drift was rejected, we reconstruct the selection (magnitude and direction) acting to diversify these groups to produce the observed differences in facial and mandibular morphology. The ancestor-descendent directionality chosen for these comparisons is consistent with our current understanding of species succession, chronology and derived versus ancestral traits. Differential selection vectors are calculated as the product of the difference vectors between fossil taxa multiplied by the inverse of the pooled within-species variance/covariance (V/CV) matrix derived from a model of *H. sapiens* variation (Table 5; see *Materials and Methods*). These vectors are visualized in Fig. 4 and Fig. 5.

**Fig. 4.**
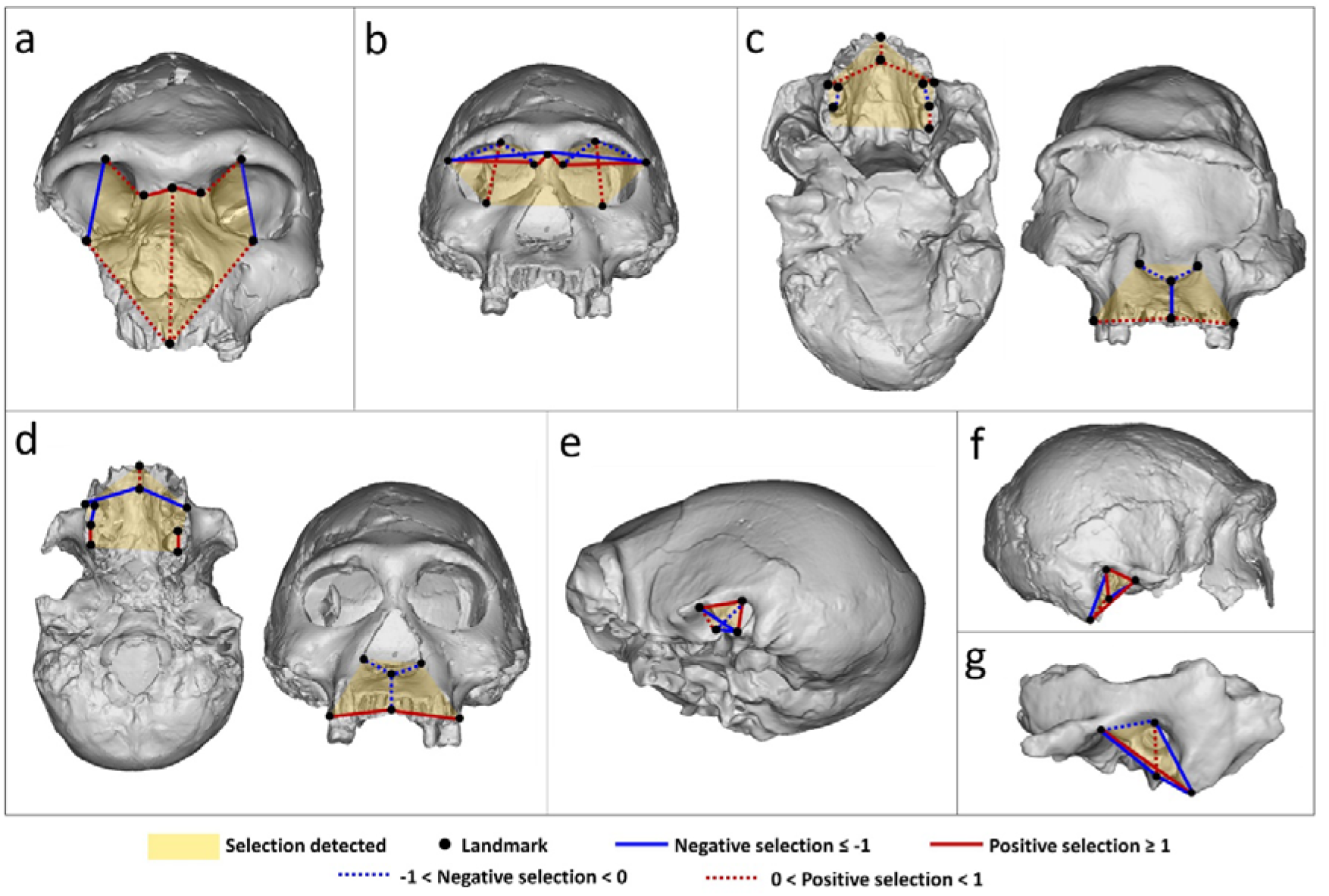
A visual representation of the selection vectors necessary to produce observed differences in cranial morphology. Landmark definitions are given in Fig. 1 and Table 1. Selection vector values are presented in Table 5. The regions undergoing selection are shaded in yellow. Positive and negative selection vectors are depicted in red and blue respectively. Strongly positive (values ≥ 1) and strongly negative (values ≤ -1) selection are represented by solid lines. Moderate to weak selection (0 > values > 1; -1 < values < 0) are displayed as dashed lines. (a) Cranial analysis 1. Selection required to produce Middle Pleistocene *Homo* from African *H. erectus* is positive for facial length/height and width and negative for superoinferior orbit height. (b) Cranial analysis 2. Selection required to produce Dmanisi *H. erectus* from *H. habilis* is positive for orbital dimensions and width of nasal bridge, and negative for upper facial width. (c) Cranial analysis 4. Selection required to produce Dmanisi *H. erectus* from A.L.666-1 is negative for maxilla height and nasal aperture width, and positive for palate depth and width. Selection on upper molar mesiodistal length varies from weakly positive to weakly negative. (d) Cranial analysis 4. Selection required to produce Dmanisi *H. erectus* from *H. habilis* is also negative for maxilla height and nasal aperture width, and positive for palate depth. Negative selection is detected for a measure of palate width. (e) Cranial analysis 6. Selection required to produce African *H. erectus* from *H. habilis* is positive for external auditory meatus (EAM) superoinferior height and mandibular fossa length. Selection is negative for the position of EAM relative to the mandibular fossa. (f) Cranial analysis 7. Selection required to produce east African *H. erectus* from Stw 53 is positive for EAM height and overall temporal shape. Negative selection affects the area between the mandibular fossa and EAM, as well as the position of the mastoid relative to porion. (g) Cranial analysis 7. Selection required to produce Middle Pleistocene *Homo* from African *H. erectus* is also positive for EAM size, however, overall temporal shape and size is mostly influenced by negative selection.

**Fig. 5.**
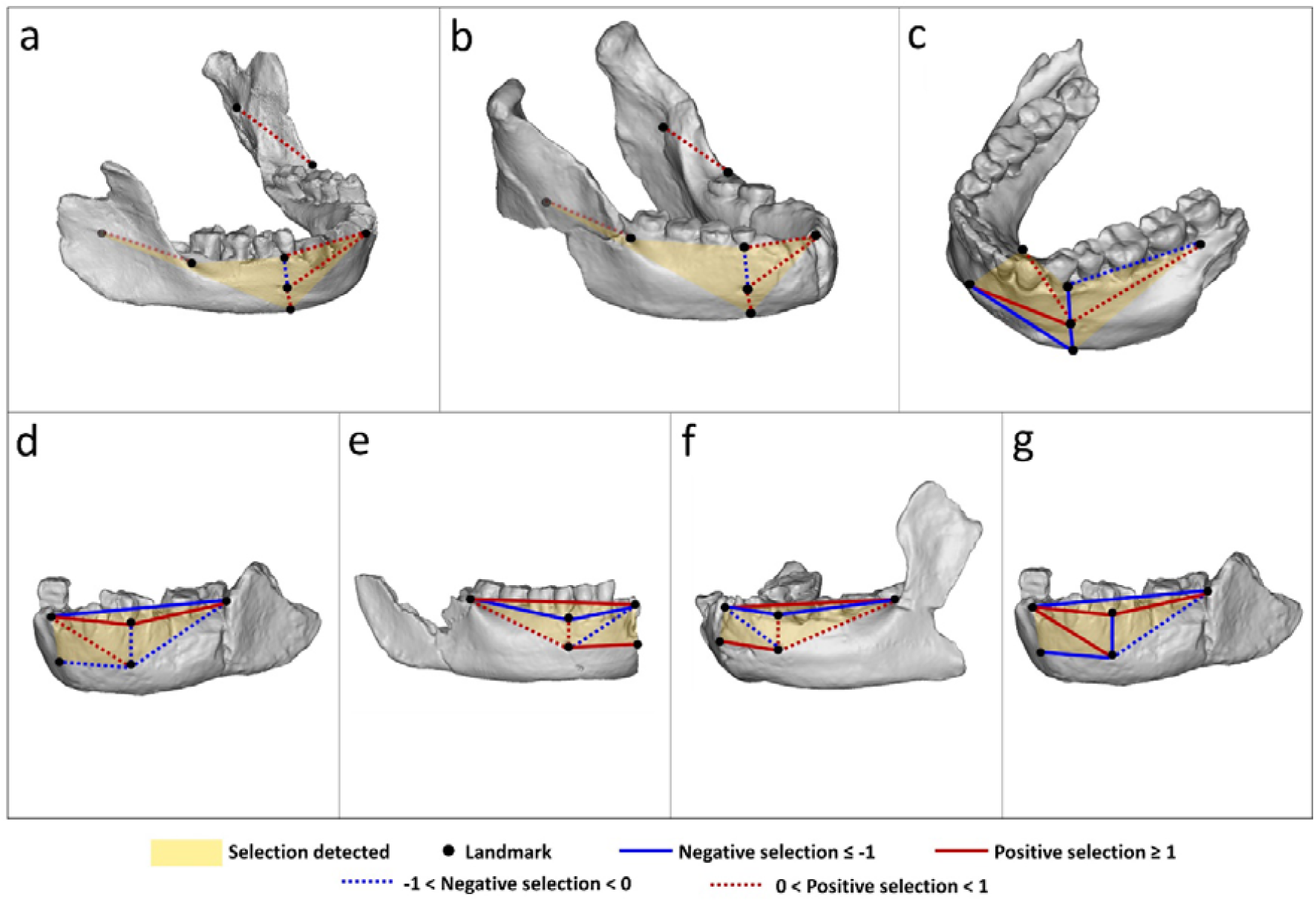
A visual representation of the selection vectors necessary to produce observed differences in mandibular morphology. Landmark definitions are given in Fig. 1 and Table 1. Selection vector values are presented in Table 5. The mandibular regions undergoing selection are shaded in yellow. Positive and negative selection vectors are depicted in red and blue respectively. Strongly positive (values ≥ 1) and strongly negative (values ≤ -1) selection are represented by solid lines. Moderate to weak selection (0 > values > 1; -1 < values < 0) are displayed as dashed lines. (a) Mandibular analysis 1. The selection required to produce Dmanisi *H. erectus* from *H. habilis* is positive for all traits, with the exception of the superoinferior position of the mental foramen which is shaped by weak negative selection. (b) Mandibular analysis 1. The selection required to produce *H. erectus (sensu lato)* from *H. habilis* displays the same pattern as the previous image. (c) Mandibular analysis 2. The selection required to produce Dmanisi *H. erectus* from *H. rudolfensis* is negative for mandibular corpus height and anterior corpus length and generally positive for posterior length and corpus thickness. (d) Mandibular analysis 4. The selection required to produce *H. rudolfensis* from *H. habilis* is mixed, acting on mandibular corpus anteroposterior length and mental foramen position. (e) Mandibular analysis 4. The selection required to produce African *H. erectus* from *H. rudolfensis* is also mixed, with selection vectors acting in the opposite direction to those seen in the previous image. (f) Mandibular analysis 4. The selection vectors required to produce *H. erectus (sensu lato)* from *H. rudolfensis* are the same as those in the previous image. (g) Mandibular analysis 4. The selection required to produce *H. rudolfensis* from LD 350-1 displays a similar pattern to that seen in (d).

**Table 5.**
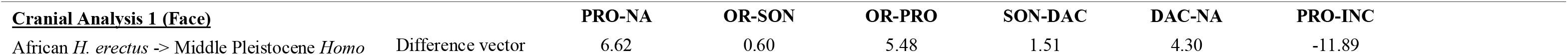
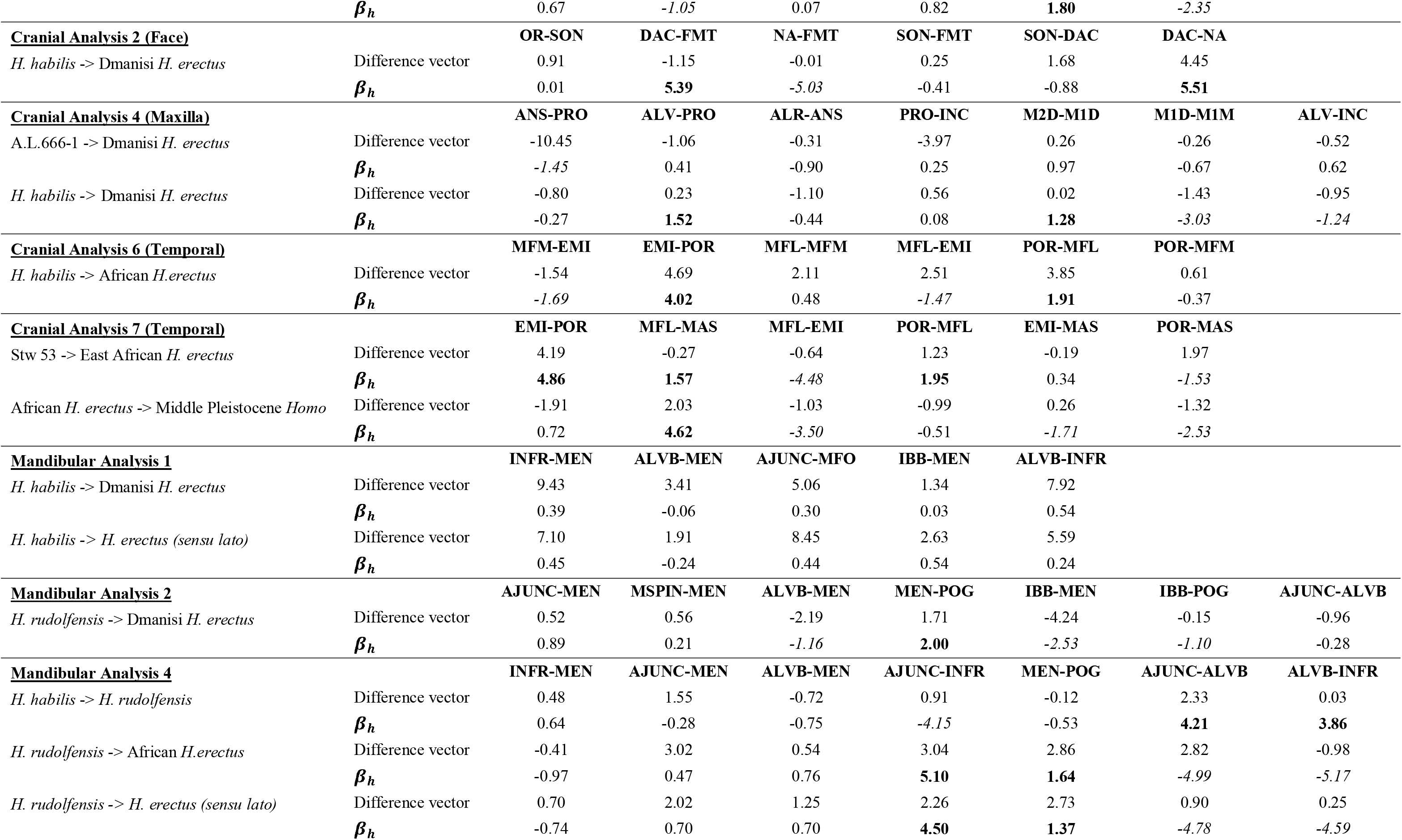
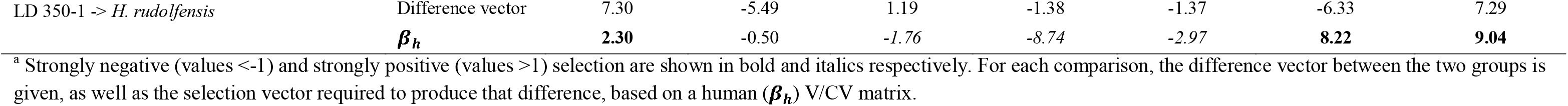
Reconstructed differential selection vectors describing the selection needed to produce later *Homo* from early *Homo*^a^

| **<u>Cranial Analysis 1 (Face)</u>** |  | **PRO-NA** | **OR-SON** | **OR-PRO** | **SON-DAC** | **DAC-NA** | **PRO-INC** |
| --- | --- | --- | --- | --- | --- | --- | --- |
| African *H. erectus* -> Middle Pleistocene *Homo* | Difference vector | 6.62 | 0.60 | 5.48 | 1.51 | 4.30 | -11.89 |

|  |  |  |  |  |  |  |  |  |
| --- | --- | --- | --- | --- | --- | --- | --- | --- |
| | $\beta_h$ | 0.67 | -1.05 | 0.07 | 0.82 | 1.80 | -2.35 | |
| **<u>Cranial Analysis 2 (Face)</u>** |  | **OR-SON** | **DAC-FMT** | **NA-FMT** | **SON-FMT** | **SON-DAC** | **DAC-NA** |  |
| *H. habilis* -> Dmanisi *H. erectus* | Difference vector | 0.91 | -1.15 | -0.01 | 0.25 | 1.68 | 4.45 |  |
| | $\beta_h$ | 0.01 | 5.39 | -5.03 | -0.41 | -0.88 | 5.51 | |
| **<u>Cranial Analysis 4 (Maxilla)</u>** |  | **ANS-PRO** | **ALV-PRO** | **ALR-ANS** | **PRO-INC** | **M2D-M1D** | **M1D-M1M** | **ALV-INC** |
| A.L.666-1 -> Dmanisi *H. erectus* | Difference vector | -10.45 | -1.06 | -0.31 | -3.97 | 0.26 | -0.26 | -0.52 |
| | $\beta_h$ | -1.45 | 0.41 | -0.90 | 0.25 | 0.97 | -0.67 | 0.62 |
| *H. habilis* -> Dmanisi *H. erectus* | Difference vector | -0.80 | 0.23 | -1.10 | 0.56 | 0.02 | -1.43 | -0.95 |
| | $\beta_h$ | -0.27 | 1.52 | -0.44 | 0.08 | 1.28 | -3.03 | -1.24 |
| **<u>Cranial Analysis 6 (Temporal)</u>** |  | **MFM-EMI** | **EMI-POR** | **MFL-MFM** | **MFL-EMI** | **POR-MFL** | **POR-MFM** |  |
| *H. habilis* -> African *H. erectus* | Difference vector | -1.54 | 4.69 | 2.11 | 2.51 | 3.85 | 0.61 |  |
| | $\beta_h$ | -1.69 | 4.02 | 0.48 | -1.47 | 1.91 | -0.37 | |
| **<u>Cranial Analysis 7 (Temporal)</u>** |  | **EMI-POR** | **MFL-MAS** | **MFL-EMI** | **POR-MFL** | **EMI-MAS** | **POR-MAS** |  |
| Stw 53 -> East African *H. erectus* | Difference vector | 4.19 | -0.27 | -0.64 | 1.23 | -0.19 | 1.97 |  |
| | $\beta_h$ | 4.86 | 1.57 | -4.48 | 1.95 | 0.34 | -1.53 | |
| African *H. erectus* -> Middle Pleistocene *Homo* | Difference vector | -1.91 | 2.03 | -1.03 | -0.99 | 0.26 | -1.32 |  |
| | $\beta_h$ | 0.72 | 4.62 | -3.50 | -0.51 | -1.71 | -2.53 | |
| **<u>Mandibular Analysis 1</u>** |  | **INFR-MEN** | **ALVB-MEN** | **AJUNC-MFO** | **IBB-MEN** | **ALVB-INFR** |  |  |
| *H. habilis* -> Dmanisi *H. erectus* | Difference vector | 9.43 | 3.41 | 5.06 | 1.34 | 7.92 |  |  |
| | $\beta_h$ | 0.39 | -0.06 | 0.30 | 0.03 | 0.54 | | |
| *H. habilis* -> *H. erectus* (sensu lato) | Difference vector | 7.10 | 1.91 | 8.45 | 2.63 | 5.59 |  |  |
| | $\beta_h$ | 0.45 | -0.24 | 0.44 | 0.54 | 0.24 | | |
| **<u>Mandibular Analysis 2</u>** |  | **AJUNC-MEN** | **MSPIN-MEN** | **ALVB-MEN** | **MEN-POG** | **IBB-MEN** | **IBB-POG** | **AJUNC-ALVB** |
| *H. rudolfensis* -> Dmanisi *H. erectus* | Difference vector | 0.52 | 0.56 | -2.19 | 1.71 | -4.24 | -0.15 | -0.96 |
| | $\beta_h$ | 0.89 | 0.21 | -1.16 | 2.00 | -2.53 | -1.10 | -0.28 |
| **<u>Mandibular Analysis 4</u>** |  | **INFR-MEN** | **AJUNC-MEN** | **ALVB-MEN** | **AJUNC-INFR** | **MEN-POG** | **AJUNC-ALVB** | **ALVB-INFR** |
| *H. habilis* -> *H. rudolfensis* | Difference vector | 0.48 | 1.55 | -0.72 | 0.91 | -0.12 | 2.33 | 0.03 |
| | $\beta_h$ | 0.64 | -0.28 | -0.75 | -4.15 | -0.53 | 4.21 | 3.86 |
| *H. rudolfensis* -> African *H. erectus* | Difference vector | -0.41 | 3.02 | 0.54 | 3.04 | 2.86 | 2.82 | -0.98 |
| | $\beta_h$ | -0.97 | 0.47 | 0.76 | 5.10 | 1.64 | -4.99 | -5.17 |
| *H. rudolfensis* -> *H. erectus* (sensu lato) | Difference vector | 0.70 | 2.02 | 1.25 | 2.26 | 2.73 | 0.90 | 0.25 |
| | $\beta_h$ | -0.74 | 0.70 | 0.70 | 4.50 | 1.37 | -4.78 | -4.59 |
| LD 350-1 -> *H. rudolfensis* | Difference vector | 7.30 | -5.49 | 1.19 | -1.38 | -1.37 | -6.33 | 7.29 |
| | $\beta_h$ | **2.30** | -0.50 | -1.76 | -8.74 | -2.97 | **8.22** | **9.04** |

<sup>a</sup> Strongly negative (values <-1) and strongly positive (values >1) selection are shown in bold and italics respectively. For each comparison, the difference vector between the two groups is given, as well as the selection vector required to produce that difference, based on a human ( $\beta_h$ ) V/CV matrix.

#### African *H. erectus* – Middle Pleistocene *Homo* (face)

The selection required to produce a Middle Pleistocene *Homo* face from an African *H. erectus* face is strongly to moderately positive for facial length, facial height and nasal bridge width, and strongly negative for superoinferior orbit height and palate depth (Fig. 4a). The response to this selection (difference vector) is mostly correlated with the direction of the selection acting on these traits, with a positive response across all variables, except palate depth. One aspect of morphology, orbit height, appears to be evolving in an opposite direction to the direction of the selection.

#### *H. habilis –* Dmanisi *H. erectus* (face)

The selection required to produce Dmanisi *H. erectus* from *H. habilis* is strongly positive to nil for the nasal bridge and orbit height and strongly to moderately negative for upper facial width and orbit width (Fig. 4b). The actual response to this selective pressure is strongly positive to nil for most variables of the upper face, except DAC-FMT, expressed as an increase in overall size of the nasal bridge and orbit in the Dmanisi hominins. For three of the variables, the response to the selective pressure is opposite to the direction of the selection, indicating that positive selection on certain traits is sufficient to drive a mostly positive response in others.

#### A.L.666-1 – Dmanisi *H. erectus* (maxilla)

The selection required to produce a Dmanisi *H. erectus* maxilla from A.L.666-1 is strongly to moderately negative for maxilla height and nasal aperture width, and moderately to weakly positive for palate depth and width (Fig. 4c). The selection on upper molar mesiodistal length varies from weakly positive to weakly negative selection. The response to selection is negative for the majority of these maxillary traits, expressed as a decrease in overall size of the maxilla. This suggests that negative selection acting on maxilla height and nasal aperture width is sufficient to drive the decrease observed in other variables.

#### *H. habilis* – Dmanisi *H. erectus* (maxilla)

The selection required to produce a Dmanisi *H. erectus* maxilla from *H. habilis* is similar to the previous instance, displaying moderately negative selection for maxilla height and nasal aperture width, and positive for palate depth (Fig. 4d). However, negative selection is detected for a measure of palate width. The morphological response resulting from these selective forces is consistent with the direction of the selection across all variables.

#### *H. habilis* – African *H. erectus* (temporal)

The selection required to produce an African *H. erectus* temporal from *H. habilis* is strongly to moderately positive for external auditory meatus (EAM) superoinferior height, POR-MFL, and mandibular fossa length (Fig. 4e). Selection is negative for the position of EAM relative to the mandibular fossa, potentially related to variability of EAM orientation. The response to this selection is positive for the majority of variables, indicating a general increase in temporal size.

#### South African early *Homo –* east African *H. erectus* (temporal)

The selection required to produce an east African *H. erectus* temporal from Stw 53 is strongly to moderately positive for EAM height and general temporal shape (Fig. 4f). Negative selection affects the area between the mandibular fossa and EAM, as well as the position of the mastoid relative to porion. The response to selection is mixed, with a positive response in EAM size and negative response in the distance between EAM and the mandibular fossa.

#### African *H. erectus –* Middle Pleistocene *Homo* (temporal)

The selection required to produce a Middle Pleistocene *Homo* temporal from African *H. erectus* is strongly to moderately positive for EAM size, as well as for the distance between the most inferior point on the mastoid process (MAS) and the mandibular fossa, however, overall temporal shape and size is mostly influenced by strongly to moderately negative selection (Fig. 4g). The response to selection is generally negative for all variables, except those including MAS. This could be indicative of an increase in the robusticity and size of the mastoid process, and could be related to sexual dimorphism.

#### *H. habilis* – Dmanisi *H. erectus* (mandible)

The selection required to produce a Dmanisi *H. erectus* mandible from *H. habilis* is moderately positive for all traits, with the exception of the superoinferior position of the mental foramen which is shaped by weak negative selection. The response to selection is positive for all traits (Fig. 5a).

#### *H. habilis* – *H. erectus* (*sensu lato*) (mandible)

The selection required to produce a *H. erectus* (*sensu lato*) mandible from *H. habilis* displays the same pattern as the previous instance (Fig. 5b).

#### *H. rudolfensis* – Dmanisi *H. erectus* (mandible)

The selection required to produce a Dmanisi *H. erectus* mandible from *H. rudolfensis* is strongly to moderately negative for mandibular corpus height, corpus length and development of pogonion, and moderately to strongly positive for posterior corpus length and corpus thickness (Fig. 5c). The direction of morphological change is consistent with the direction of the selection pressures, expressed as an increase in overall corpus length and thickness and a decrease in corpus height. It also suggests possible selective pressure on the position of the mental foramen (MEN).

#### *H. habilis – H. rudolfensis* (mandible)

The selection required to produce a *H. rudolfensis* mandible from *H. habilis* is moderately to strongly positive for mandibular corpus anteroposterior length, and moderately to strongly negative for the relative position of MEN, as well as for traits describing the shape of the mandibular arcade (Fig. 5d). The response to selection is mostly positive, with the exception of traits describing the position of MEN.

#### *H. rudolfensis* – African *H. erectus* (mandible)

The pattern of selection required to produce an African *H. erectus* mandible from *H. rudolfensis* is similar to the previous instance, except completely opposite in direction. Strongly negative selection is detected for mandibular corpus anteroposterior length, and moderately to strongly positive selection for the relative position of MEN, as well as for traits describing the shape of the mandibular arcade (Fig. 5e). The response to selection is mostly positive, with the exception of two traits relating to the relative anteroposterior position of MEN and anterior corpus length. The direction of morphological change is consistent with the direction of selection pressure for most traits, except AJUNC-ALVB, which increases despite negative selection.

#### *H. rudolfensis* – *H. erectus* (*sensu lato*) (mandible)

The pattern of selection required to produce a *H. erectus* <u>(*sensu lato*)</u> mandible from *H. rudolfensis* is the same as the previous comparison (Fig. 5f). The response to selection is positive for all traits.

#### LD 350-1 – *H. rudolfensis* (mandible)

The selection required to produce a *H. rudolfensis* mandible from LD 350-1 displays a similar pattern to that seen in (Fig. 5d), where selection is strongly positive for mandibular corpus anteroposterior length, and moderately to strongly negative for the relative position of MEN, as well as for traits describing the shape of the mandibular arcade (Fig. 5g). The response to selection is mixed, with a positive response for traits associated with the position of MEN and/or corpus height, and a negative response for traits describing corpus length. This is expressed as an increase in corpus length and a decrease in corpus height and/or a more superior location of MEN.

## Discussion

The results of our analyses indicate that morphological relationships among *Homo* taxa are complex, and suggest that diversification may be driven primarily (though not exclusively) by neutral evolution. Multivariate and geometric morphometric results were generally consistent and highlighted the large amount of morphological diversity within *Homo,* especially within *H. erectus,* a geographically and temporally widespread species. Other interesting patterns also emerged. First, the spatial relationships among specimens differed depending on the morphological region analyzed. For example, Mahalanobis’ distances between *H. erectus* specimen KNM-ER 3883 and other Pleistocene *Homo* are significantly different for the temporal region (Fig. 2c), but not for the face (Fig. 2a) and neurocranium (Fig. 2d). Second, the Dmanisi hominins and specimens of *H. rudolfensis* are consistently different from each other and from other taxa. Third, the oldest *Homo* specimen, LD 350-1, is significantly different from all other specimens for calculations of Mahalanobis’ distances, except for *H. erectus* specimen KNM-BK 8518 and *H. sapiens* specimen Tuinplaas 1. This specimen also falls within, or on the boundary of, the *H. erectus* convex hulls in principal component plots of Procrustes shape coordinates (Figs 3a-b), lending support to the initial diagnosis of this specimen as *Homo* (Villmoare et al., 2015). Finally, it is worth noting that there is a close association between *H. naledi* and *H. erectus* in both cranial and mandibular analyses (e.g. similar to what has been shown in Dembo et al., 2016; Laird et al., 2016; Schroeder et al., 2016), as well as between ∼2.4 Ma early *Homo* specimen A.L.666-1, South African specimen Stw 53, and *H. habilis* specimen KNM-ER 1813. The results of these metric analyses confirm the complexity of the phenotypic variation within *Homo* and the difficulty faced when trying to identify potential evolutionary relationships, especially given the possibility multiple lineages within our genus.

What has produced this diversity? Our results indicate that for 95% of taxon comparisons (51% when a conservative estimate of statistical power is used), across the entire skull (face, maxilla, neurocranium, temporal, mandible), the null hypothesis of genetic drift cannot be rejected. This indicates that of the majority of the cranial and mandibular phenotypic diversity within *Homo,* from ∼2.8 Ma-0.0117 Ma, is consistent with random genetic drift. This is particularly striking for the neurocranium where all three analyses comprising 39 different comparisons are shown to be consistent with drift, even when including very small-brained *H. erectus* (Dmanisi) and *H. naledi* (South Africa). What this indicates is that the relative size and shape variation that exists between taxa is proportional to that seen within taxa (here based on the *Homo sapiens* model). In other words, although morphological divergence is occurring among species, it happens consistently across the phenotype in a manner that does not change the relative relationships among parts. For the neurocranium, this is true despite considerable brain size differences between *Homo* taxa. In this light, recent suggestions that brain size and shape differences may poorly define *Homo* (Spoor et al., 2015) are intriguing, because they have arisen in the context of an increased understanding of comparable magnitudes and patterns of variation within taxa. It may be more difficult to delineate taxa under a model of drift, as opposed to a model of selection, which drives changes in the relative relationships among traits. However, it is important to remember that the neurocranial analyses in particular, due to a dearth of available homologous landmarks, did not capture all aspects of brain shape but rather gross shape/size. Nonetheless, based on these results it is necessary to re-consider the traditional view that selection was the main evolutionary process driving changes in the neurocranium, and most other cranial regions, within *Homo*, and consider the implications of that for our understanding of how and why our lineage evolved.

For the remaining cases, where drift was rejected, three primary patterns can be observed. First, adaptation played a role in driving the evolution of differences between the Dmanisi hominins and other early *Homo* specimens across both the face and mandible. Interestingly, even though the Dmanisi group itself is hugely diverse, we found that this rejection of drift is consistent across all of the Dmanisi specimens, regardless of the specimen or combination of specimens included in each analysis, confirming that this result was not just a product of intra-group variability. The Dmanisi hominins were the first of our lineage to leave Africa, and our results indicate that selection played an important role in that dispersal, resulting in significant morphological changes (and a different covariance structure) as these hominins adapted to new environmental contexts. Second, although drift was the primary force implicated in neurocranial change, selection repeatedly acted to shape maxillary and mandibular diversity among *Homo* groups. This result suggests that the evolution of *Homo* is characterized by adaptive diversification in masticatory systems among taxa, which may be related to dietary change, possibly as a result of environmental change (Vrba, 1985, 1995, 1996, 2007; Cerling, 1992; Stanley, 1992; deMenocal, 1995; Reed, 1997; Bobe and Behrensmeyer, 2004; Wynn, 2004), environment variability (Potts, 1998), and/or shifts to new foraging strategies (Stanley, 1992; Braun et al., 2010; Lepre et al., 2011; Potts, 2012; Ferraro et al., 2013). Third, the mandibular morphology of *H. rudolfensis* consistently emerges as being adaptively different from other *Homo* taxa, including the earliest *Homo* specimen, LD 350-1. This result implies a potentially divergent and distinct evolutionary trajectory for this taxon, possibly signifying a branching event, supporting the distinctiveness of this taxon, and providing an adaptive explanation for divergence in sympatry with other *Homo* taxa (i.e. *H. habilis*). However, despite these instances where drift was rejected, we reiterate that, for the majority, selection was not detected. For some cases, this lack of selection is surprising. For example, we do not see a massive adaptive change occurring between 2.7 and 2.5 Ma as per Vrba’s 1985 turnover-pulse hypothesis (Vrba, 1985), nor do we see the expected correspondence between most major cultural transitions and changes in skull morphology.

Interestingly, we also do not detect major selective pressure acting to differentiate *Homo sapiens* from Middle Pleistocene *Homo.* This result parallels the findings of Weaver et al. 2007 who show that genetic drift can account for the cranial differences between Neanderthals and modern humans. It also provides further evidence for a “lengthy process model” of modern human origins (Weaver, 2012), supporting the theory of morphological continuity from the later Middle Pleistocene, ∼400 000 years ago, to the appearance of anatomically modern humans. While it is important to note that these analyses were only performed on crania and mandibles, these results are nonetheless significant given the emphasis placed on cranial and mandibular material for alpha taxonomy.

There is a fundamental disconnection between the realization that molecular change over evolutionary timeframes occurs predominantly through neutral processes (Kimura, 1968, 1991), and the dominant interpretation (explicitly or implicitly) that morphological change in human evolution is primarily adaptive and directional. The results of this study lend further support to the notion that random change has played a major role in human evolution (see also Ackermann and Cheverud, 2004; Weaver et al., 2007; Schroeder et al., 2014). The detection of widespread genetic drift acting on all aspects of skull morphology during the evolution of our genus is likely to be due, in part, to small population sizes of groups in isolation. This could also be correlated with a purported population bottleneck at ∼2.0 Ma (Hawks et al., 2000). Because the emergence and evolution of *Homo* and the appearance and proliferation of stone tools roughly correspond, and continue to co-evolve, it is also possible that hominins were increasingly reliant on cultural adaptations – as opposed to biological adaptations – to manage environmental changes (Schroeder et al., 2014; Ackermann and Cheverud, 2004; Lynch, 1990). Continued investigation into evolutionary process is necessary – especially for anatomical regions such as the postcranium which remain largely unexplored (but see Grabowski and Roseman, 2015) – in order to provide further insight into how and why the human lineage evolved.

## Acknowledgements

We would like to thank the many funding agencies that supported this research: Palaeontological Scientific Trust (L.S.), the L.S.B. Leakey Foundation (Baldwin Fellowship; L.S.), the South African National Research Foundation (L.S., R.R.A.), and the DST/NRF Centre of Excellence in Palaeosciences (COE-Pal; R.R.A.). We are grateful to the many institutions and individuals for access to comparative fossil and extant primate and human materials in their care: E. Mbua, P. Kiura, and the National Museums of Kenya; the National Museum and House of Culture of Tanzania; National Museum of Ethiopia; S. Potze and the Ditsong Museum; B. Billings and the School of Anatomical Sciences of the University of the Witwatersrand; B. Zipfel and the Evolutionary Studies Institute at the University of the Witwatersrand; W. Seconna of the Iziko South African Museum; Y. Haile-Selassie and L.M. Jellema of the Cleveland Museum of Natural History; and F. Schrenk, O. Kullmer, and C. Hemm at the Senckenberg Forschungsinstitut. We also thank K. Reed and W. Kimbel for granting access to a 3-D scan of LD 350-1.

## References

Ackermann, R.R., 1998. A quantitative assessment of variability in the australopithecine, human, chimpanzee, and gorilla face. Ph.D. Dissertation, Washington University.

Ackermann, R.R., 2003. Using extant morphological variation to understand fossil relationships: a cautionary tale. S. Afr. J. Sci. 99, 255–258.

Ackermann, R.R., Cheverud, J.M., 2004. Detecting genetic drift versus selection in human evolution. Proc. Natl. Acad. Sci. USA 101, 17946–17951.

Antón, S.C., Potts, R., Aiello, L.C., 2014. Evolution of early *Homo*: An integrated biological perspective. Science 345, 1236828.

Berger, L.R., Hawkes, J., de Ruiter, D.J., Churchill, S.E., Schmid, P., Williams, S.A., DeSilva, J.M., Kivell, T.L., Skinner, M.M., Musiba, C.M., Cameron, N., Holliday, T.W., Harcourt-Smith, W.E.H., Ackermann, R.R., Bastir, M., Bogin, B., Bolter, D.R., Brophy, J.K., Cofran, Z.D., Congdon, K.A., Deane, A.S., Delezene, L.K., Dembo, M., Drapeau, M., Elliott, M.E., Feuerriegel, E.M., Garcia-Martinez, D., Garvin, H.M., Green, D.J., Gurtov, A.N., Irish, J.D., Kruger, A., Laird, M.F., Marchi, D., Meyer, M.R., Nalla, S., Negash, E.W., Orr, C.M., Radovcic, D., Schroeder, L., Scott, J.E., Throckmorton, Z., Trocheri, M.W., VanSickle, C., Walker, C.S., Wei, P., Zipfel, B., 2015. A new species of *Homo* from the Dinaledi Chamber, South Africa. Elife 4, e0956.

Bobe, R. and Behrensmeyer, A.K., 2004. The expansion of grassland ecosystems in Africa in relation to mammalian evolution and the origin of the genus *Homo*. Palaeogeogr. Palaeoclimatol. Palaeoecol. 207, 399–420.

Braun, D.R., Harris, J.W., Levin, N.E., McCoy, J.T., Herries, A.I., Bamford, M.K., Bishop, L.C., Richmond, B.G., Kibunjia, M., 2010. Early hominin diet included diverse terrestrial and aquatic animals 1.95 Ma in East Turkana, Kenya. Proc. Natl. Acad. Sci. USA 107, 10002–10007.

Brown, P., Sutikna, T., Morwood, M.J., Soejono, R.P., Saptomo, E.W., Due, R.A., 2004. A new small-bodied hominin from the Late Pleistocene of Flores, Indonesia. Nature 431, 1055–1061.

Cerling, T.E., 1992. Development of grasslands and savannas in East Africa during the Neogene. Palaeogeogr. Palaeoclimatol. Palaeoecol. 97, 241–247.

Champely, S., 2016. pwr: Basic Functions for Power Analysis. R package version 1.2-0. http://CRAN.R-project.org/package=pwr.

Cheverud, J.M., 1988. A comparison of genetic and phenotypic correlations. Evolution 42, 958-968.

Dembo, M., Radovcic, D., Garvin, H.M., Laird, M.F., Schroeder, L., Scott, J.E., Brophy, J., Ackermann, R.R., Musiba, C.M., de Ruiter, D.J., Mooers, A. Ø., Collard, M. 2016. The evolutionary relationships and age of *Homo naledi*: An assessment using dated Bayesian phylogenetic methods. J. Hum. Evol. 97, 17–26.

Dirks, P.H., Roberts, E.M., Hilbert-Wolf, H., Kramers, J.D., Hawks, J., Dosseto, A., Duval, M., Elliot, M., Evans, M., Grün, R., Hellstrom, J., Herries, A.I.R, Joannes-Boyau, R., Makhubela, T.V., Placzek, C.J., Robbins, J., Spandler, C., Wiersma, J., Woodhead, J., Berger, L.R. 2017. The age of *Homo naledi* and associated sediments in the Rising Star Cave, South Africa. eLife 6, e24231

deMenocal, P.B., 1995. Plio-Pleistocene African climate. Science 270, 53–59.

Ferraro, J.V., Plummer, T.W., Pobiner, B.L., Oliver, J.S., Bishop, L.C., Braun, D.R., Ditchfield, P.W., Seaman III, J.W., Binetti, K.M., Seaman Jr, J.W., Hertel, F., 2013. Earliest archaeological evidence of persistent hominin carnivory. PLoS one 8(4), e62174.

Gower, J.C., 1975. Generalized procrustes analysis. Psychometrika 40, 33–51.

Grabowski, M., Roseman, C.C., 2015. Complex and changing patterns of natural selection explain the evolution of the human hip. J. Hum. Evol. 85, 94–110.

Hammer, Ø., Harper, D.A.T., Ryan, P.D., 2001. PAST: Paleontological Statistics Software Package for Education and Data Analysis. Palaeontol. Electron. 4, 1–9.

Harmand, S., Lewis, J., Feibel, C.S., Lepre, C.J., Sandrine, P., Lenoble, A., Boës, X., Quinn, R., Brenet, M., Arroyo, A., Taylor, N., Clément, S., Daver, G., Brugal, J.-P., Leakey, L., Mortlock, R.A., Wright, J.D., Lokorodi, S., Kirwa, C., Kent, D.V., Roche, H., 2015. 3.3- million-year-old stone tools from Lomekwi 3, West Turkana, Kenya. Nature 521, 310–315.

Harvati, K., 2003. The Neanderthal taxonomic position: models of intra-and inter-specific craniofacial variation. J. Hum. Evol. 44, 107–132.

Hawks, J., Hunley, K., Lee, S. H., Wolpoff, M., 2000. Population bottlenecks and Pleistocene human evolution. Mol. Biol. Evol. 17, 2–22.

Kimura, M., 1968. Evolutionary rate at the molecular level. Nature 217, 624–626.

Kimura, M., 1991. The neutral theory of molecular evolution: a review of recent evidence. Jpn. J. Genet. 66, 367–386.

Klingenberg, C.P., 1996. Multivariate allometry. In: Marcus, L.F., Corti, M., Loy, A., Naylor, G.J.P., Slice, D.E. (Eds.), Advances in Morphometrics. Plenum, New York, pp. 23–49.

Klingenberg, C.P., 1998. Heterochrony and allometry: the analysis of evolutionary change in ontogeny. Biol. Rev. 73, 70–123.

Klingenberg, C.P., 2011. MorphoJ: an integrated software package for geometric morphometrics. Mol. Ecol. Resour. 11, 353–357.

Klingenberg, C.P., 2016. Size, shape, and form: concepts of allometry in geometric morphometrics. Dev. Genes Evol. 226, 113–137.

Kramer, A., Donnelly, S., Kidder, J., Ousley, D., Olah, S.M., 1995. Craniometric variation in large-bodied hominoids: testing the single species hypothesis for *Homo habilis*. J. Hum. Evol. 29, 443–462.

Laird, M.F., Schroeder, L., Garvin, H.M., Scott, J.E., Dembo, M., Radovcic, D., Musiba, C.M., Ackermann, R.R., Schmid, P., Hawks, J., Berger, L.R., de Ruiter, D.J., 2016. The skull of Homo naledi. J. Hum. Evol. In press. DOI: 10.1016/j.jhevol.2016.09.009.

Lande, R., 1977. Statistical tests for natural selection on quantitative characters. Evolution 31, 442–444.

Lande, R., 1979. Quantitative genetic analysis of multivariate evolution, applied to brain: body size allometry. Evolution 33, 402–416.

Lande, R., 1980. Genetic variation and phenotypic evolution during allopatric speciation. Am. Nat. 116, 463–479.

Lande, R., Arnold, S.J., 1983. The measurement of selection on correlated characters. Evolution 37, 1210–1226.

Leakey, L.S.B., Tobias, P.V., Napier, J.R., 1964. A new species of the genus *Homo* from Olduvai Gorge. Nature 202, 7–9.

Lepre, C.J., Roche, H., Kent, D.V., Harmand, S., Quinn, R.L., Brugal, J.P., Texier, P.J., Lenoble, A., Feibel, C.S., 2011. An earlier origin for the Acheulian. Nature 477(7362), 82–85.

Lieberman, D.E., Wood, B.A., Pilbeam, D.R., 1996. Homoplasy and early *Homo*: an analysis of the evolutionary relationships of *H. habilis sensu stricto* and *H. rudolfensis*. J. Hum. Evol. 30, 97–120.

Lockwood, C.A., Kimbel, W.H., Lynch, J.M., 2004. Morphometrics and hominoid phylogeny: support for a chimpanzee-human clade and differentiation among great ape subspecies. Proc. Natl. Acad. Sci. 101, 4356–4360.

Lordkipanidze, D., Ponce de León, M.S., Margvelashvili, A., Rak, Y., Rightmire, G.P., Vekua, A., Zollikofer, C.P.E., 2013. A complete skull from Dmanisi, Georgia, and the evolutionary biology of early *Homo*. Science 342, 326–331.

Lynch, M., 1990. The rate of morphological evolution in mammals from the standpoint of the neutral expectation. Am. Nat. 136, 727–741.

Miller, J.A., 1991. Does brain size variability provide evidence of multiple species in *Homo habilis*? Am. J. Phys. Anthropol. 84, 385–398.

Miller, J.A., 2000. Craniofacial variation in *Homo habilis*: an analysis of the evidence for multiple species. Am. J. Anthropol. 112, 103–128.

Mitteroecker, P., Gunz, P., Windhager, S., Schaefer, K., 2003. A brief review of shape, form, and allometry in geometric morphometrics, with applications to human facial morphology. Hystrix 24, 59–66.

Monteiro, L.R., 1999. Multivariate regression models and geometric morphometrics: The search for causal factors in the analysis of shape. Syst. Biol. 48, 192–199.

Potts, R. 2012. Environmental and behavioral evidence pertaining to the evolution of early Homo. Curr. Anthropol. 53, S299–S317.

Potts, R., 1998. Variability selection in hominid evolution. Evol. Anthropol. 7, 81–96.

Reed, K.E., 1997. Early hominid evolution and ecological change through the African Plio-Pleistocene. J. Hum. Evol. 32, 289–322.

Rohlf, F.J., 1999. Shape statistics: Procrustes superimpositions and tangent spaces. J. Classif. 16, 197–223.

Rohlf, F.J., Slice, D.E., 1990. Extensions of the Procrustes method for the optimal superimposition of landmarks. Syst. Biol. 39, 40–59.

Schroeder, L., Roseman, C.C., Cheverud, J.M., Ackermann, R.R., 2014. Characterizing the Evolutionary Path(s) to Early *Homo*. PLoS one 9, e114307.

Schroeder, L., Scott, J.E., Garvin, H.M., Laird, M.F., Dembo, M., Radovcic, D., Berger, L.R., de Ruiter, D.J., Ackermann, R.R., 2016. Skull diversity in the *Homo* lineage and the relative position of *Homo naledi*. J. Hum. Evol. In press.

Smith, H. F., 2009. Which cranial regions reflect molecular distances reliably in humans? Evidence from three-dimensional morphology. Am. J. Hum. Biol. 21, 36–47.

Spoor, F., Gunz, P., Neubauer, S., Stelzer, S., Scott, N., Kwekason, A., Dean, M.C., 2015. Reconstructed *Homo habilis* type OH 7 suggests deep-rooted species diversity in early *Homo*. Nature 519(7541), 83–86.

Stanley, S.M., 1992. An ecological theory for the origin of *Homo*. Paleobiol. 18, 237–257.

Villmoare, B., Kimbel, W.H., Seyoum, C., Campisano, C.J., DiMaggio, E.N., Rowan, J., Braun, D.R., Arrowsmith, J.R., Reed, K.E., 2015. Early *Homo* at 2.8 Ma from Ledi-Geraru, Afar, Ethiopia. Science 347, 1352–1355.

von Cramon-Taubadel, N., 2009. Revisiting the homoiology hypothesis: the impact of phenotypic plasticity on the reconstruction of human population history from craniometric data. J. Hum. Evol. 57, 179–190.

Vrba, E.S., 1985. Ecological and adaptive changes associated with early hominid evolution. In: Delson E. (Ed.), Ancestors: The Hard Evidence. A.R. Liss, New York, pp. 63–71.

Vrba, E.S., 1995. On the connections between paleoclimate and evolution. In: Vrba E.S., Denton G.H., Partridge T.C., Burckle L.H. (Eds.), Paleoclimate and Evolution with Emphasis on Human Origins. Yale University Press, New Haven, CT, pp. 24–45.

Vrba, E.S., 1996. Climate, heterochrony, and human evolution. J. Anthropol. Res. 52, 1-28.

Vrba, E.S., 2007. Role of environmental stimuli in hominid origins. In: Henke W., Rothe H., Tattersall I. (Eds.), Handbook of Paleoanthropology: Phylogeny of Hominines. Springer, Berlin, Heidelberg, pp. 1441–1481.

Weaver, T.D., 2012. Did a discrete event 200 000–100 000 years ago produce modern humans? J. Hum. Evol. 63, 121–126.

Weaver, T.D., Roseman, C.C., Stringer, C.B., 2007. Were neandertal and modern human cranial differences produced by natural selection or genetic drift? J. Hum. Evol. 53, 135–145.

Williams, F.L., Richtsmeier, J.T., 2003. Comparison of mandibular landmarks from computed tomography and 3D digitizer data. Clin. Anat. 16, 494–500.

Willmore, K.E., Roseman, C.C., Rogers, J., Cheverud, J.M., Richtsmeier, J.T., 2009. Comparison of mandibular phenotypic and genetic integration between baboon and mouse. Evol. Biol. 36, 19–36.

Wood, B., 1992. Origin and evolution of the genus *Homo*. Nature 355, 783–790.

Wood, B., 1993. Early *Homo*: How many species. In: Kimbel W.H., Martin L.B. (Eds.), Species, Species Concepts, and Primate Evolution. Plenum Press, New York, pp. 485–522.

Wood, B., Baker, J., 2011. Evolution in the genus Homo. Annu. Rev. Ecol. Evol. Syst. 42, 47–69.

Wynn, J.G., 2004. Influence of Plio-Pleistocene aridification on human evolution: Evidence from paleosols of the Turkana Basin, Kenya. Am. J. Phys. Anthropol. 123, 106–118.

